# The competitive interplay of 12-oxophytodienoic acid (OPDA), protein thiols and glutathione

**DOI:** 10.1101/2025.07.31.667886

**Authors:** Madita Knieper, Ruben Schwarz, Lara Vogelsang, Jens Sproß, Armağan Kaya, Maike Bittmann, Harald Gröger, Andrea Viehhauser, Karl-Josef Dietz

## Abstract

*Cis*-(+)-12-Oxophytodienoic acid (OPDA) is a bioactive oxylipin and phytohormone participating in regulation of plant stress responses, growth and development. Due to its α,β-unsaturated carbonyl moiety, OPDA covalently binds to free thiol groups by Michael addition. This binding, termed OPDAylation, alters the activity of target proteins, such as cyclophilin 20-3 (EC:5.2.1.8) and thioredoxins, that are essential components of the cellular redox regulatory network. To function as reversible redox regulatory mechanism, OPDAylation should be complemented by a process of de-OPDAylation allowing for fine-tuning of OPDA-dependent regulation. This study explored OPDAylation and de-OPDAylation *in vitro* with emphasis on involvement of glutathione. OPDA can be transferred from protein to glutathione (GSH), and *vice versa*. In a competition experiment, OPDAylation of thioredoxins occurred rapidly in the presence of GSH, while over extended incubation times, de-OPDAylation of TRX occurred due to the stoichiometric excess of GSH. These results support the hypothesis that the initial thioredoxin-based OPDAylation is proceeding under kinetic control due to the higher reactivity of the more nucleophilic cysteine moiety in thioredoxin compared to the one of GSH, while the OPDAylation of GSH observed at prolonged incubation time is then a result of a thermodynamically controlled process. De-OPDAylation depends on the protein’s sensitivity towards OPDA, the pH and the concentration of excess thiol groups. This likely allows for precise modulation of OPDA amounts, as the rapid modification of protein activity enables subsequent induction of OPDA signaling, whereas de-OPDAylation, triggered by increasing glutathione, increasing cellular reduction or presumably enzymatically, reverses this effect.

## 1. Introduction

*Cis*-(+)-12-oxophytodienoic acid (OPDA) is a cyclopentenone oxylipin, which is synthesized in the chloroplast from α-linolenic acid by 13-lipoxygenase (13-LOX, EC:1.13.11.12), allene oxide synthase (AOS, EC:4.2.1.92) and allene oxide cyclase (AOC, EC:5.3.99.6) [1]. While it is a precursor for the plant hormone jasmonic acid (JA), OPDA itself acts as a phytohormone affecting crucial processes such as germination, plant growth and stress defense [2–6].

OPDA bioactivity originates from its cyclopentenone moiety which assigns it to the group of reactive carbonyl species (RCS) as highly reactive α,β-unsaturated carbonyl compound that forms covalent adducts with free thiol groups by Michael addition [7– 9]. Formation of protein OPDA adducts by Michael addition, termed OPDAylation, was previously described as a novel posttranslational modification (PTM) affecting thiol switch proteins like thioredoxins and, thereby, modulating the redox regulatory network of plant cells [10–12].

The most prominent target of OPDAylation in *Arabidopsis* is cyclophilin 20-3 (Cyp20-3). Formation of Cyp20-3-OPDA adducts enhances formation of the cysteine synthase complex (CSC), activating thiol synthesis and subsequently increasing the cellular reduction potential. This increase, in turn, modulates the expression of OPDA-responsive genes (ORGs) that contribute e.g. to plant defense under wounding or high light stress [11, 13].

High light stress is a frequently occurring abiotic stress condition that is initiated if the photosynthetic active radiation exceeds the growth light, and in full sunlight reaching >2000 µmol quanta m^-2^ s^-1^ in temperate latitudes [14, 15]. Excess excitation energy (EEE) enhances the production of reactive oxygen species (ROS) such as H_2_O_2_ and O_2.-_ in the chloroplast. To cope with EEE, defense mechanisms are needed to adjust ROS scavenging and maintain cellular redox balance [16, 17]. In this context, adjustment of gene expression occurs within seconds [18]. Recent studies positioned OPDA as the responsible jasmonate coordinating expression of stress defense genes under early-stage EEE stress with JA-mediating plant recovery only afterwards [17, 19, 20]. Moreover, OPDA-specific signaling presumably also extends to long-term high light stress, as indicated by the uncoupling of OPDA and JA synthesis [21, 3]. At this point, OPDA-mediated photoprotection traces back to enhanced ROS scavenging and chaperone capacities as well as synthesis of non-photosynthetic pigments like flavonoids and glucosinolates [20, 17, 11, 22–24, 19].

Nevertheless, as RCS like OPDA accumulate, they can cause oxidative damage to proteins, lipids and DNA [7, 25–27, 22]. Therefore, tight regulation of OPDA levels is essential. Recently, conjugates of OPDA and various amino acids have been discovered in *A. thaliana* [28–30] and rice [31]. In contrast to JA-isoleucine, these conjugates, like dn-OPDA-amino acid-conjugates [32], might be inactive compounds contributing to the homeostasis of OPDA levels as OPDA storage pools [30].

Furthermore, detoxification of OPDA occurs by adduct formation (GS-OPDA) with glutathione (GSH), the major low molecular mass thiol antioxidant in eukaryotes, and subsequent vacuolar degradation (catalyzed by γ-glutamyl transpeptidase 4 [GGT4, EC:2.3.2.2]) [27, 33]. Various glutathione-*S*-transferases (GSTs) of the phi and tau class have been attributed to GS-OPDA formation, with GST activities towards OPDA ranging from 0.1-0.3 nkat mg^-1^ (GSTU1, GSTU2, GSTU4, GSTU5) and 0.4-0.8 nkat mg^-1^ (GSTU8, GSTU19) to 1.0-1.6 nkat mg^-1^ (GSTF8, GSTU6, GSTU10, GSTU17, GSTU25) [34, 35].

In general, to efficiently participate in dynamic regulation, reversibility of a particular PTM is an important prerequisite. Although Michael addition is widely regarded as a relatively stable process with low reversibility [36], reversibility of RCS coupling contributes to fine-tuning of RCS signaling as has been shown for 4-HNE, acrolein, prostaglandins and isoprostanes [37–41].

Since recovery of OPDAylated proteins has not been studied so far, this investigation was designed to explore whether OPDA-thiol adducts might undergo *retro*-Michael addition, allowing for (i) improved fine-tuning of OPDA signaling under stress conditions and (ii) potential recovery of free OPDA after adduct formation.

The latter would be of special interest as it might explain how a free OPDA pool for OPDA signaling and conversion to JA or OPDA-amino acid-adducts is maintained in the presence of high chloroplastic and cytosolic thiol levels.

Furthermore, this study gives an insight into the competing roles of cysteine-containing moieties in the formation of OPDAylated Michael-type products, exemplified for thioredoxin and GSH as *S*-nucleophiles.

Our work addresses this dilemma *in vitro* by employing thioredoxins and Cyp20-3 as selected target proteins of OPDAylation. Additionally, H_2_O_2_ and flavonoid levels of *A. thaliana* leaf discs exposed to EEE grant insight to OPDA signaling and its possible reversibility *in vivo*. The results reveal a dynamic network of OPDA, protein-OPDA adducts and GS-OPDA, hypothetically dependent on variation of GST expression.

## 2. Material and Methods

### 2.1 Expression and Purification of Recombinant Proteins

Recombinant His-tagged proteins were expressed in *E.coli* NiCo 21 (DE3) cells containing plasmids encoding for At3g62030 (Cyp20-3), At1g63460 (GPXL8), At2g06050 (OPR3, EC:1.3.1.42), At2g17420 (NTRA, EC:1.8.1.9), At5g42980 (TRX-h3), At3g02730 (TRX-f1) and At3g15360 (TRX-m4), without transit peptides as described in [42], [10, 43], [44] and [12]. Expression was induced by the addition of 400 µM isopropyl-β-D-thiogalactopyranoside in the exponential phase and incubation at 37°C for 4 h. For OPR3 expression, 20 µg/l riboflavin was added and expression period was prolonged to 20 h at room temperature (RT). Cells were harvested by centrifugation at 11000 *xg* at 4°C for 10 min, or, in case of Cyp20-3 and OPR3, at 4500 *xg* and 4°C for 15 or 30 min, respectively.

After purification and dialysis, the purity of the proteins was assessed by Coomassie staining after separation in 12% (w/v) SDS-PAGE, and a Bradford assay (Roti®Quant, Roth, Karlsruhe, Germany) was conducted to determine the protein concentration.

### 2.2 Synthesis of OPDA

12-OPDA was provided and synthesized as described in previous literature. The spectroscopic data of the product was in accordance with the published data [45, 46].

### 2.3 OPDAylation of thiol groups

Prior to OPDA treatment, 100 µM purified proteins were reduced by 10 mM DTT (RT, 1 h) and desalted using a PD10 column according to the manufacturer’s instructions (gravity protocol, Merck, Darmstadt, Germany). 20 µM reduced protein was then incubated with indicated concentrations of OPDA or equal concentrations of EtOH as a control (RT, 16 h, if not specified otherwise).

### 2.4 Mass-spectrometric analysis of OPDAylated thiols

#### 2.4.1 MS analysis of protein thiols

MS analysis was performed by applying 10 µM protein (in 50% ACN+ FA) to a Q-IMS-TOF mass spectrometer Synapt G2Si (Waters GmbH, Manchester, UK) operated in resolution mode and interfaced to a nano-ESI ion source. N_2_ generated with the nitrogen generator NGM 11 served as nebulizer and dry gas. Argon was used as collision gas for MS2 experiments using collision induced dissociation (CID). Samples were introduced by static nano-ESI using in-house pulled glass emitters. External calibration of the mass axis used ESI-L Tuning Mix (Agilent Technologies, Santa Clara, CA, USA) as standard. Scan accumulation and data processing was performed with MassLynx 4.1 (Waters GmbH, Manchester, UK) on a PC workstation. The final spectra were generated by accumulating and averaging 50 single spectra. Determination of exact masses of OPDA was performed based on centroided data from the protonated signal of leucine-enkephalin as an internal mass standard. Determination of protein masses was performed with the tool of Winkler [22]. Prior to protein mass determination, MS data were baseline subtracted, smoothed and centroided.

To test different OPDA:TRX ratios, TRX-h3 was prepared as specified above and eluted in 50 mM ammonium acetate. 20 µM reduced TRX-h3 was treated with 20-80 µM OPDA (or 0.07-0.27 % [v/v]) EtOH overnight and then subjected to MS analysis. For stability analysis of OPDAylated TRX-h3, 20 µM TRX-h3 was incubated with 80 µM OPDA overnight, then desalted (Zeba Spin Desalting Columns 7K MWCO, Thermo Scientific, Darmstadt, Germany) to remove excess OPDA and applied to MS after 0, 3, 6 or 24 h. De-OPDAylation assays of TRX-h3 were performed in the same manner, however after desalting, 1 mM GSH was added to the sample.

#### 2.4.2 MS analysis of non-protein thiols

Mass spectrometry was performed analogously to 2.4.1, with minor adjustments: 150 µM OPDA were incubated with 1 mM GSH or DTT (in 50 mM ammonium acetate) overnight and either analyzed directly, or, for competition assays, 1 mM thiol compounds as indicated was added, and presence of the OPDA adducts was analyzed before, and directly after addition, as well as after 1, 2, 4 or 24 h.

### 2.5 Quantification of free thiol groups

250 µM of DTT or GSH were incubated with 500 µM OPDA (or 0.56% [v/v] EtOH as control) in 40 mM KPi buffer of different pH values (pH 5.7-8.7) for 30 min at 750 rpm shaking speed and 25°C. Residual free DTT or GSH was determined by 5,5’-dithiobis-2-nitrobenzoic acid (DTNB) assay using a standard curve of 0-500 µm GSH [12]. Samples were measured at 412 nm with a plate reader (KC4, BIOTEK Instruments).

### 2.6 Insulin reduction assay

TRX-mediated insulin reduction was performed as described in [10]. Briefly, 20 µM TRX-h3, TRX-f1 or TRX-m4 were incubated with 80 µM OPDA or EtOH (0.27% [v/v] as control) at RT for 1 h with subsequent addition of 1 mM GSH or equal volume of water as a control. To study the effect of GSH pretreatment, protein was first incubated with GSH or water followed by addition of OPDA or equal EtOH concentrations as control. To measure insulin precipitation (in a total volume of 200 µl), 5 µM of treated protein was added to 100 mM KPi, pH 7.2, 1 mM EDTA and 160 µM insulin in a 96-well plate. After measuring baseline absorbance, DTT (final concentration of 1 mM) was added, and absorbance was recorded at 650 nm for 60 min.

### 2.7 TRX-dependent peroxidase activity

TRX activity was measured as electron donor to glutathione peroxidase-like 8 (GPXL8) after addition of H_2_O_2_ according to [10]. Briefly, 20 µM of reduced TRX-h3 was incubated with 150 µM OPDA (or 0.5% [v/v] EtOH) for 1 h at RT. Next, 0.25-5 mM GSH was added and 0.25 µM TRX was added to a reaction mix containing 200 µM NADPH, 1 µM NADPH-dependent thioredoxin reductase A (NTRA) and 3 µM GPXL8 (in 0.1 M Tris-HCl, pH 7.5, 5 mM EDTA). After recording the baseline for 3 min, addition of 300 µM H_2_O_2_ started the enzyme assay. The consumption of NADPH+H^+^ by NTRA was monitored at 340 nm (Cary 3500 UV-Vis, Agilent, Santa Clara, CA, USA).

### 2.8 Cyp20-3 activity

PPIase activity of Cyp20-3 was measured spectrophotometrically as described in [12]. Reduced Cyp20-3 (5, 10, 25, 50, 75, 100 nM) was mixed with 100 µM N-succinyl-Ala-Ala-Pro-Phe-p-nitroanilide and either 25 µM OPDA, 0.083% [v/v] EtOH, 1 mM DTT or 1 mM H_2_O_2_ in 35 mM HEPES, pH 8.0 in polystyrene cuvettes (total volume of 2 ml). The sample mix was incubated at 8°C and 800 rpm for 10 min. After recording the baseline at 390 nm for 3 min (Cary 3500 UV-Vis, Agilent, Santa Clara, CA, USA), the reaction was started by adding 0.4 mg/ml α-chymotrypsin. De-OPDAylation of Cyp20-3 by GSH was analyzed by first incubating 20 µM of reduced Cyp20-3 with 150 µM OPDA (50 mM HEPES, pH 8, RT, 2 h), followed by addition of 1 mM of GSH and further incubation (48 h at RT). For pH-dependency, incubation conditions were modified to 40 mM KPi adjusted to pH 6.2, 6.7, 7.2, 7.7 or 8.2 and 4 h of incubation at RT.

### 2.9 Tryptophan fluorescence

15 µM of reduced TRX-f1 or TRX-m4 (dissolved in 50 mM KPi, pH 7.2) was incubated with 150 µM OPDA, 0.5% [v/v] EtOH, 1 mM H_2_O_2_ or 1 mM DTT for 3 h. For de-OPDAylation, samples were further incubated for 16 h after addition of 1 mM GSH or DTT. To assess the influence of GSH incubation prior to OPDAylation, 15 µM reduced protein was first incubated with 1 mM GSH for 2 h, followed by addition of 150 µM OPDA (or 0.5% [v/v] EtOH as a control) and 3 h of incubation. All steps were performed at RT.

Relative tryptophan fluorescence was determined using a spectrofluorometer (SFM 25, Kontron Instruments), set to an excitation wavelength of 280 nm, a scan rate of 1 nm sec^-1^, high voltage 450 V and an emission wavelength range of 300-400 nm.

### 2.10 HPLC-based quantification of OPDA

Quantification of OPDA was performed as described in [10]. 200 µM OPDA was incubated with 50 µM TRX-f1 in 40 mM KPi pH 7.2 at RT for 2 h. 25 - 250 µM DTT was added and the residual amount of free OPDA was measured after 4 h by applying 25 µl of sample to a Dionex HPLC system equipped with a LiChrospherR 100 RP-18 column (LiChrom-CARTR 250-4, Merck, Darmstadt, Germany) set to 30°C and 1.75 mL min^-1^. After 9 min of equilibration with solution A (80% [v/v] MeOH, 0.1% [v/v] acetic acid), solution B (99.9% [v/v] MeOH, 0.1% [v/v] acetic acid) was gradually added (9-14 min 0-25% solution B, 14-16 min 25-100% solution B). Each run was recorded at 224 nm (Chromeleon Version 6.6, Dionex) for 18 min. A standard curve of 0-250 µM OPDA was used to calculate the concentration of OPDA in the samples.

### 2.11 Plant growth and preparation of leaf discs

*A. thaliana* Col-0 plants were grown in soil in a greenhouse with a day/night cycle of 8h/16h with 100 µmol photons m^-2^ s^-1^, 22°C/20°C and 45-55% relative humidity. Prior to that, seeds were incubated at 4°C for 3 d. After 35-40 d of growth, leaf discs of 5 mm diameter were cut out using a cork borer and immediately placed upside down in a 96-well-plate filled with either 25 µM oxylipin solution or, as a mock control, 0.083% (v/v) EtOH. After 16 h of incubation, 1 mM GSH (or H_2_O as a control) was added to the floating solution, leaf discs were subjected to EEE stress (800 µmol photons m^-2^ s^-1^) for 6 h and frozen in liquid nitrogen for determination of H_2_O_2_ and flavonoid levels. To analyze OPDAylation effects after GSH pretreatment, leaf discs were first incubated with 1 mM GSH for 16 hours and 25 µM OPDA (or 0.083% [v/v] EtOH) was added directly before stress treatment.

### 2.12 Measurement of H_2_O_2_ content

H_2_O_2_ content of *A. thaliana* leaf discs was determined according to [47] with minor modifications. 35-45 mg finely ground plant material (8 leaf discs shock-frozen in N_2 liquid_) was homogenized in 0.5 ml of 5% (v/v) trichloro acetic acid at 4°C and then incubated in the dark at 4°C for 40 min. To remove proteins, samples were centrifuged (10000 *xg*, 15 min, 4°C) and 50 µl of the supernatant was diluted 250-fold in 0.1 M sodium carbonate buffer, pH 10.2. Then, 20 µl of dilution was mixed with either 5 µl 50 U catalase (EC:1.11.1.6) or 5 µl H_2_O and incubated in the dark at 30°C for 15 min. Chemiluminescence was measured at 425 nm for 5-8 s after adding 2 µl of sample (catalase-treated or untreated) to 998 µl working solution (65 µM luminol, 30 µM CoCl_2_ in 0.1 M sodium carbonate buffer, pH 10.2). A standard curve was generated with H_2_O_2_ (0-15 µM).

### 2.13 Measurement of flavonoid content

Flavonoid content of leaf discs was determined based on [48] with several adjustments. 100 µl of plant extract (prepared as described in 2.12) were mixed with 400 µl of H_2_O and 30 µl of 5% (w/v) NaNO_2_ and incubated at RT for 6 min. 30 µl of 10% [w/v] AlCl_3_ were added and the reaction was stopped after 6 min by adding 400 µl 1 M NaOH. The volume was adjusted to 1 ml with H_2_O. After 15 min, 150 µl of sample was transferred to a microtiter plate and OD_405 nm_ was measured using a plate reader. Flavonoid content was calculated based on a standard curve of 0.5-10 µg quercetin.

### 2.14 Statistical analysis

To determine significant differences, either one way-ANOVA in combination with post hoc Tukey test or t-test with Bonferroni correction were carried out using RStudio (2025.05.0+496) [49].

## 3. Results

OPDA signaling function has been experimentally linked to a variety of stress defense mechanisms in plants. However, so far it remained unknown whether OPDAylation is a permanent protein modification and how it can be achieved despite high prevalence of GSH under physiological conditions. To this end, the aim of this work was to explore the effect of excess free thiols on the formation of OPDA adducts as well as the reversibility of OPDAylation (de-OPDAylation).

### 3.1 Formation of OPDA adducts

As a first step, generation of OPDA adducts was studied under control conditions to confirm the reactivity of OPDA towards thiol groups. TRX-h3 was chosen as a target protein since it is highly susceptible to OPDAylation [10]. When testing various stoichiometric ratios, a 4:1 OPDA:TRX-h3 ratio yielded fully OPDAylated protein after 16 h of incubation (Fig. 1, Table S1). Hence, this ratio was adopted for MS analyses of OPDAylated TRX.

**Figure 1:**
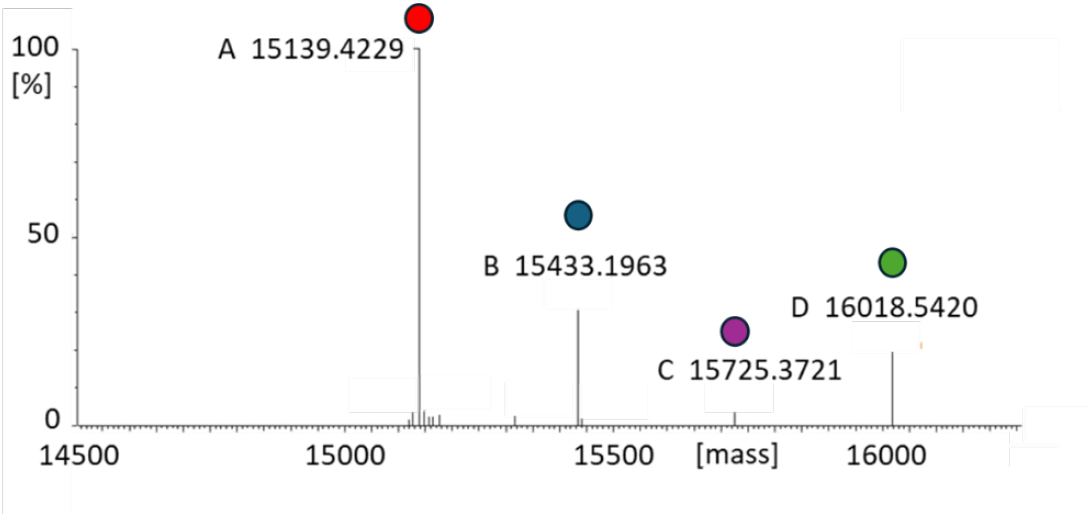
MS analysis of adduct formation between TRX-h3 and OPDA. 20 µM protein was incubated with 80 µM of OPDA in 50 mM ammonium acetate for 16 h prior to measurement. Different charge states of the protein could be detected revealing free (native) TRX (15.14 kDa, red dot) and 1x (15.43 kDa, blue dot), 2x (15.73 kDa, purple dot) and 3x OPDAylated protein (16.02 kDa, green dot).

Binding of OPDA to non-protein thiols, namely GSH and DTT, could be observed by decrease of free thiol groups as well as MS analysis (Fig. 2). Furthermore, thiol quantification revealed a pH-dependent adduct formation, presumably by altering the protonation state of the thiol group (Fig. 2A, C). Additional analysis of GS-OPDA formation by means of ^1^H-NMR-spectroscopic studies confirmed a conversion of 59% over an incubation time of 3 h (calculated on the basis of the ^1^H-NMR-signals of the C*<u>H</u>*-C(O)-proton at 7.73 ppm and the two C*<u>H</u>*=C*<u>H</u>*-protons at 5.25 ppm) (Fig. S1).

**Figure 2:**
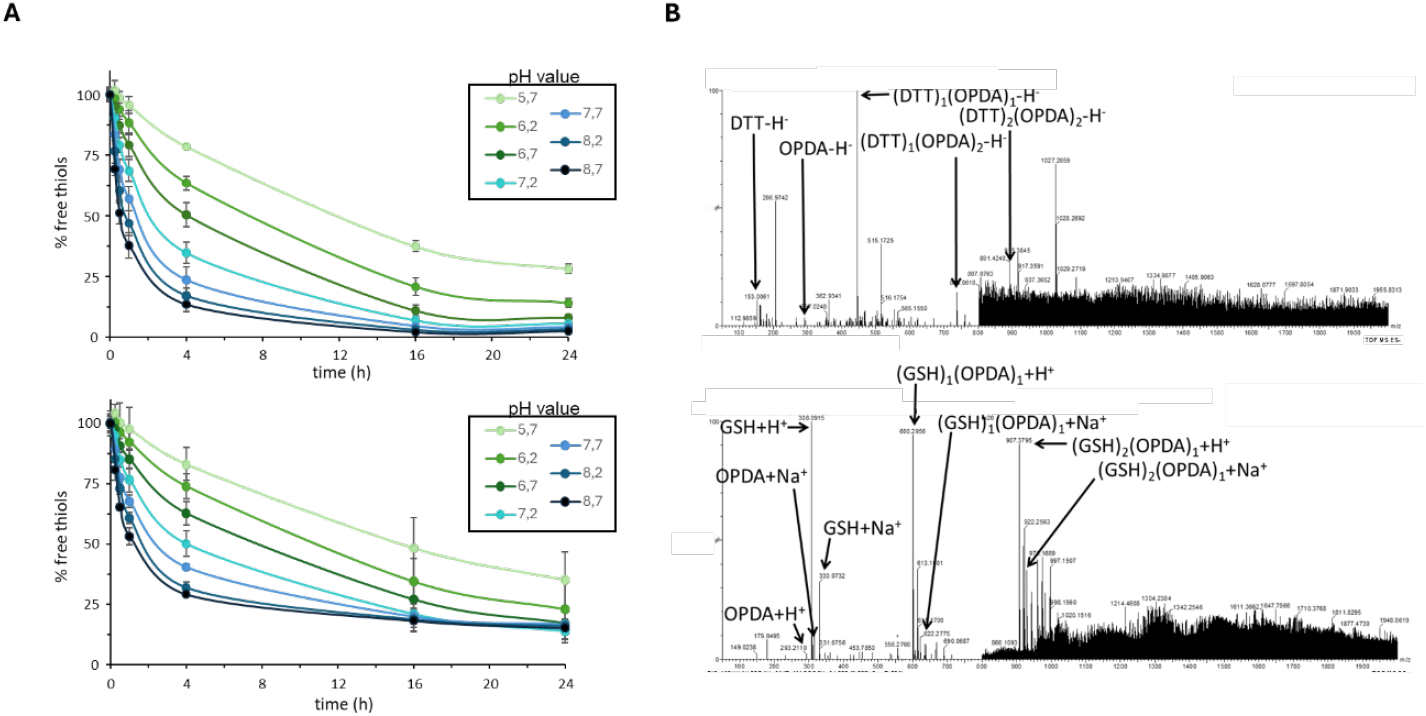
Formation of DTT or GS-OPDA adducts. OPDA (500 µM) was incubated with 250 µM DTT (A, B) or GSH (C, D) in 40 mM KPi, pH 5.7-8.7, and residual free thiols were quantified at indicated time points. Data are means ± SD of n=3. (A, C). Alternatively, 150 µM OPDA was incubated with 1 mM GSH/DTT over night and subjected to MS (B, D). Labelled arrows indicate the masses of specified molecular species and adducts of DTT and OPDA and glutathione and OPDA.

### 3.2 Disintegration of protein adducts by excess thiols

OPDA adducts proved to be stable under control conditions (Table S2). As next step, GSH was added to OPDAylated TRX-h3, and possible reversibility of OPDAylation was assessed by MS analysis. GSH was chosen as potential trigger of de-OPDAylation as GSH is the dominant low molecular mass thiol antioxidant in the cell and was previously reported to support *retro*-Michael addition of TRX-acrolein adducts [38]. Obtained spectra, resembling those depicted in Fig. 1, showed a slow gradual decrease of TRX-OPDA adducts after addition of GSH (Fig. 3, Fig. S3). The peaks corresponding to onefold, twofold and threefold OPDAylated protein vanished almost completely after 24, 30 and 50 h, respectively. OPDAylated protein was undetectable after 96 h.

**Figure 3:**
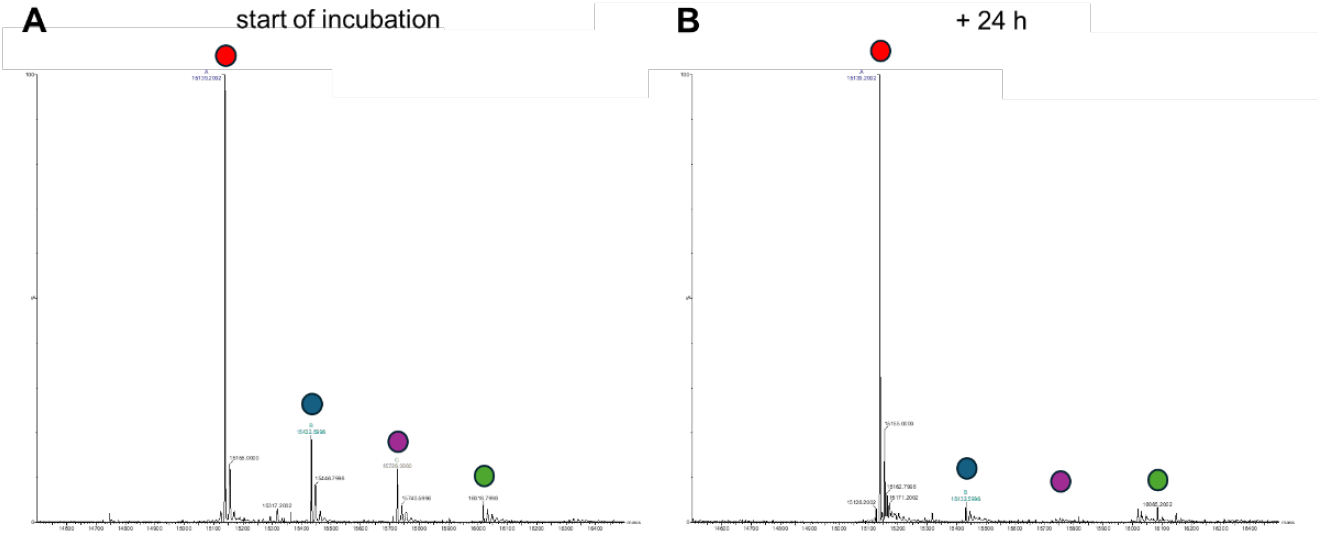
Time-dependent de-OPDAylation of TRX-h3 after addition of GSH. Recombinant TRX was reduced, desalted, treated with OPDA for 16 h and desalted again. Samples were subjected to MS analysis after addition of GSH at 0-96 h. Peaks correspond to free TRX (15.14 kDa, red dot) and 1-3x OPDAylated TRX (15,43 kDa and blue dot, 15,73 kDa and purple dot, 16,02 kDa and green dot, respectively). For MS spectra of additional timepoints and reference of OPDAylated TRX-h3 incubated without GSH see Figure S3 and Table S2.

The observed gradual decrease of TRX-OPDA adducts hinted towards a reversibility of OPDAylation by excess free thiols, however, the oxidation state and/or activity of released protein remained unclear. To address this question, protein activity was measured analogously to previous studies [11]. The insulin reduction capacity of different thioredoxins (TRX-h3, -f1 and -m4) was chosen as a general readout for disulfide reduction activity (Fig. 4) and TRX-dependent GPXL8 activity as specific reduction assay (Fig. 5). OPDAylation showed an inhibitory effect on all tested thioredoxins. The residual activity was 47% for TRX-h3, 67% for TRX-f1 and 71% for TRX-m4. Both the insulin assay and the GPXL8 assay revealed recovery of the reductase activity after addition of 1 mM GSH (Figs. 4-5).

**Figure 4:**
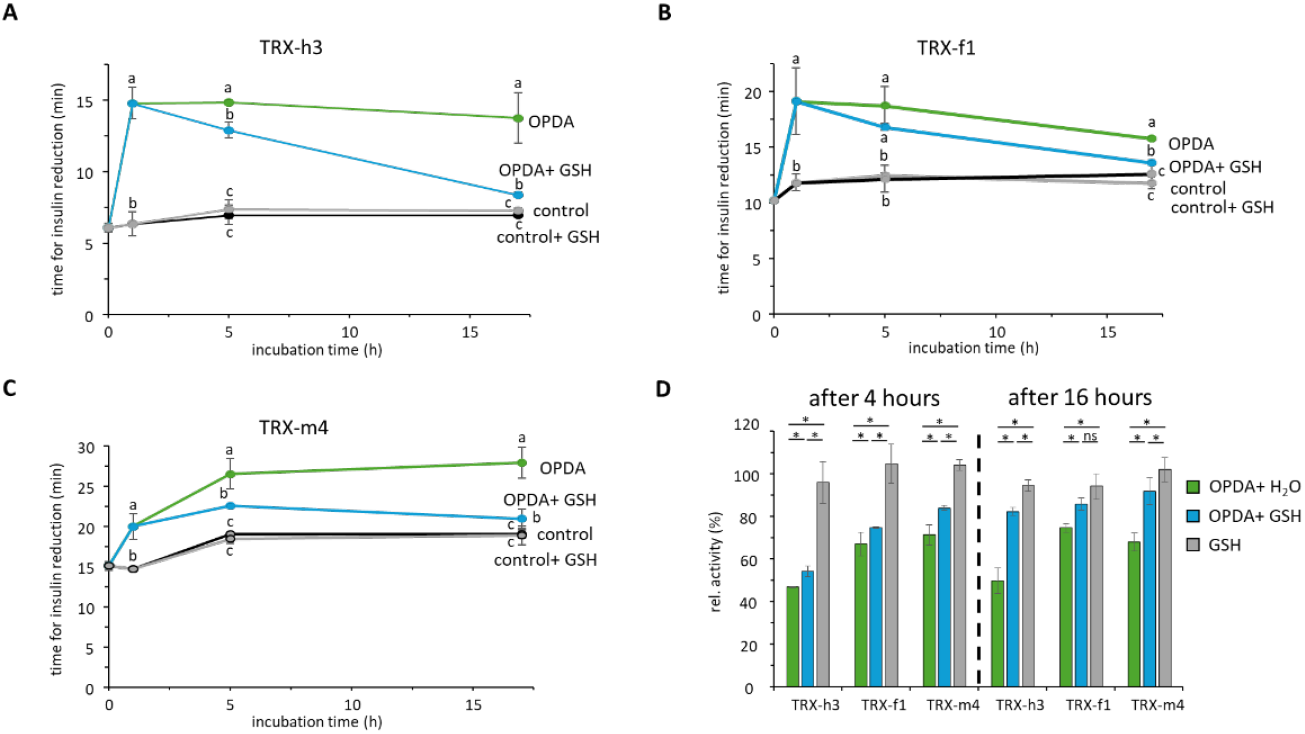
Insulin reduction activity of thioredoxins after (de-)OPDAylation. Recombinant TRX (20 µM) was treated with OPDA (80 µM) or 0.27% (v/v) EtOH as mock control for 1 h. Then, GSH, or water as mock control, was added and activity was determined after 4 and 16 h using TRX at a final concentration of 5 µM. (A-C) Protein activity is depicted as the time needed to pass the threshold of ΔA_650 nm_= 0.3. Data are means ± SD of n≥3. (D) TRX activity was determined as the change in A_650 nm_ min^-1^ and normalized to the control (mock-treated sample, 100% activity equals ΔA_650nm_ min^-1^=0.045, 0.025 and 0.016 (t4) and 0.045, 0.027 and 0.016 (t16) for TRX-h3, -f1 and -m4, respectively. Significance of difference was assessed by t-test (p<0.05) and is either denoted as letters a-c or as *.

**Figure 5:**
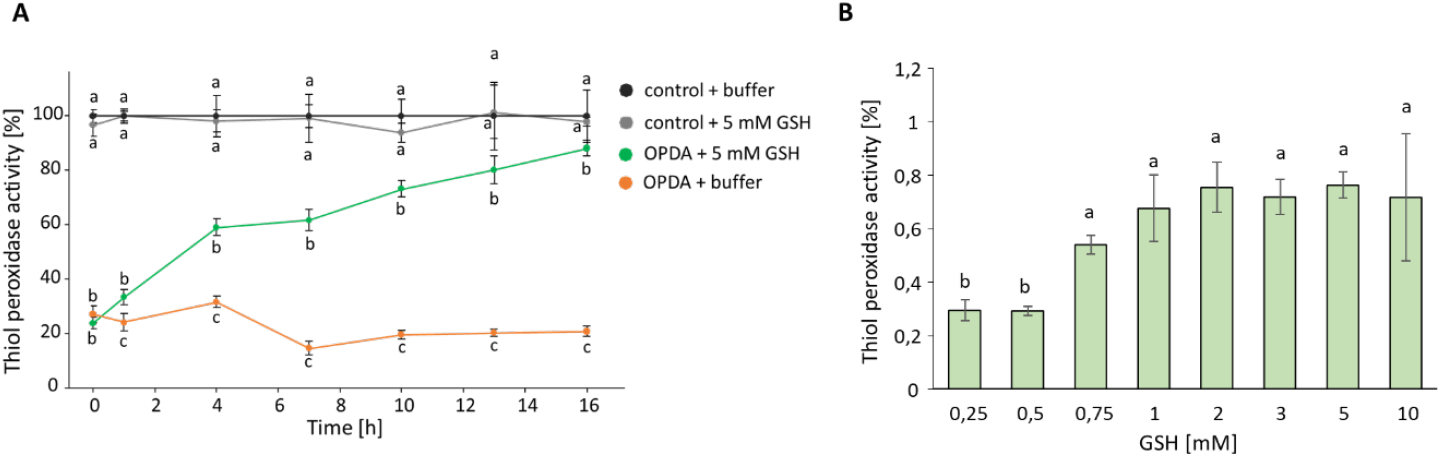
TRX-dependent H_2_O_2_ reduction activity of GPXL8. Following OPDAylation or mock treatment (0.5% [v/v] EtOH) of TRX-h3 for 1 h, GSH, or an equivalent amount of water as control, was added and GPXL8 activity with TRX-h3 as electron donor was determined using the NADPH→NTRA→TRX-h3→GPXL8→H_2_O_2_ reconstitution assay. Mean and SD for n≥3 are shown. Significances of difference between the four different samples at a given time point were determined using ANOVA, followed by posthoc Tukey test and are marked with letters a-c.

To determine the protein redox state after de-OPDAylation, intrinsic tryptophan fluorescence of TRX-f1 and TRX-m4 was measured (Table 1). After OPDAylation, relative fluorescence intensity of both proteins was quenched (−14.5 and -16.3 for TRX-f1 and -m4, respectively) to a similar level as oxidized protein. This decrease was more pronounced after longer incubation. After addition of GSH, fluorescence intensity of both TRXs increased and almost reached the value of reduced protein. It is concluded that TRX-f1 and -m4 return to the reduced state after the OPDA release. This would be in accordance with the return of protein activity as thioredoxins are redox-sensitive proteins.

**Table 1:** Intrinsic protein fluorescence of 15 µM TRX-f1 and TRX-m4 incubated with OPDA +/-GSH. Given values refer to maximal fluorescence intensity (r. U). Data are means ± SD of n=5. Statistically significant differences (t-test, p<0.05) are indicated by *.

| Incubation time | Treatment | TRX-f1 | TRX-m4 |
| --- | --- | --- | --- |
| 3 h | Control | 116.8 ± 1.9 | 115.3 ± 3.1 |
| | OPDA | 102.3 ± 1.8 * | 99.0 ± 1.7 * |
| | H<sub>2</sub>O<sub>2</sub> | 99.9 ± 1.3 * | 98.2 ± 1.9 * |
| 24 h | Control + H<sub>2</sub>O | 118.6 ± 3.1 | 118.7 ± 3.0 |
| | Control + GSH | 114.8 ± 3.5 | 117.9 ± 2.9 |
| | OPDA + H<sub>2</sub>O | 87.7 ± 2.9 * | 91.6 ± 3.3 * |
| | OPDA + GSH | 106.2 ± 3.1 * | 105.5 ± 2.5 * |

Binding of OPDA to Cyp20-3 stimulates thiol synthesis by stabilizing the Cys synthase complex. This effect is a central mechanism of OPDA-specific signaling [11]. Therefore, the next experiments addressed OPDAylation of Cyp20-3. After OPDA treatment for 10 min, PPIase activity of Cyp20-3 decreased drastically (Fig. 6A, B), which is in accordance with previous studies [10, 50]. As expected, the addition of DTT and GSH caused a partial return of catalytic efficiency (211% and 149% recovery). Like GS-OPDA formation (Fig. 2C), the degree of reactivation depended on the pH value of the medium with maximal recovery at pH 6.2 and 8.2 (Fig. 6C).

**Figure 6:**
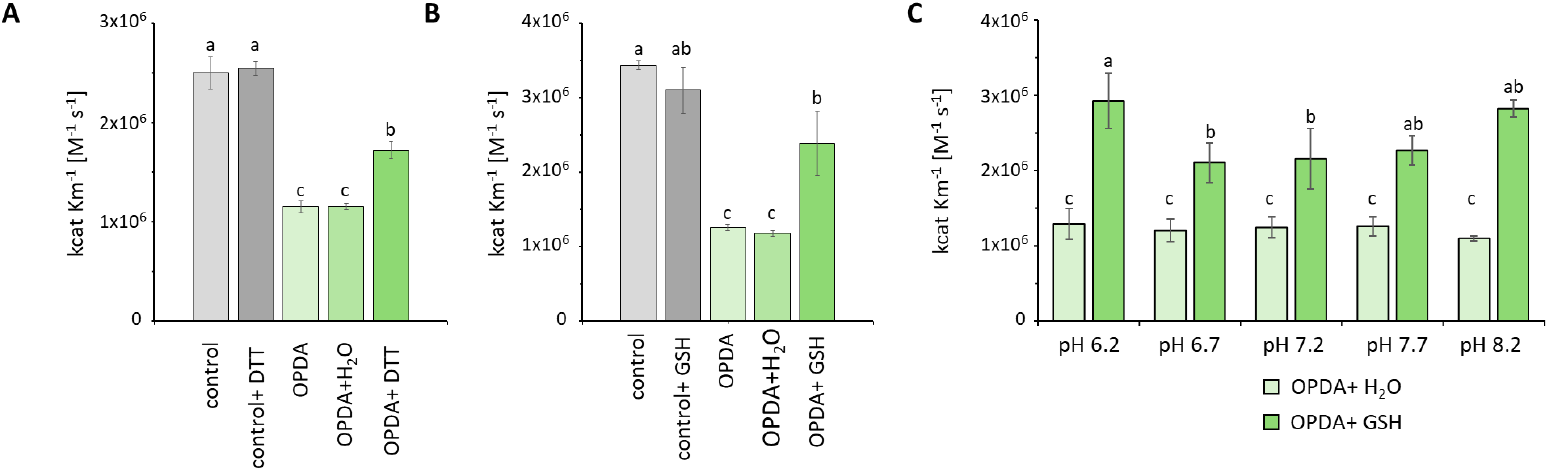
Recovery of PPIase activity of OPDAylated Cyp20-3 by GSH or DTT. Pretreatment of 20 µM Cyp20-3 with 150 µM OPDA inhibited PPIase activity. This inhibition was partially reversed upon incubation in the presence of 1 mM GSH or DTT as determined after 10 min (A), 48 h (B) or 4 hours (C) of incubation. Data are means ± SD of n≥3. Significant differences (p<0.05, marked as a-c) were calculated by one way ANOVA and Tukey test.

Lastly, release of OPDA from Michael adducts was studied by incubating GS-OPDA with DTT (Fig. 7A) and DTT-OPDA with GSH (Fig. 7B) and subsequent determination of OPDA adducts by MS. As expected, both DTT and GSH caused disintegration of adducts while forming new conjugates to OPDA. Interestingly, addition of GSH to DTT-OPDA adducts caused complete vanishing of DTT-adducts after 24 hours, while GS-OPDA adducts remained stable in the presence of DTT.

**Figure 7:**
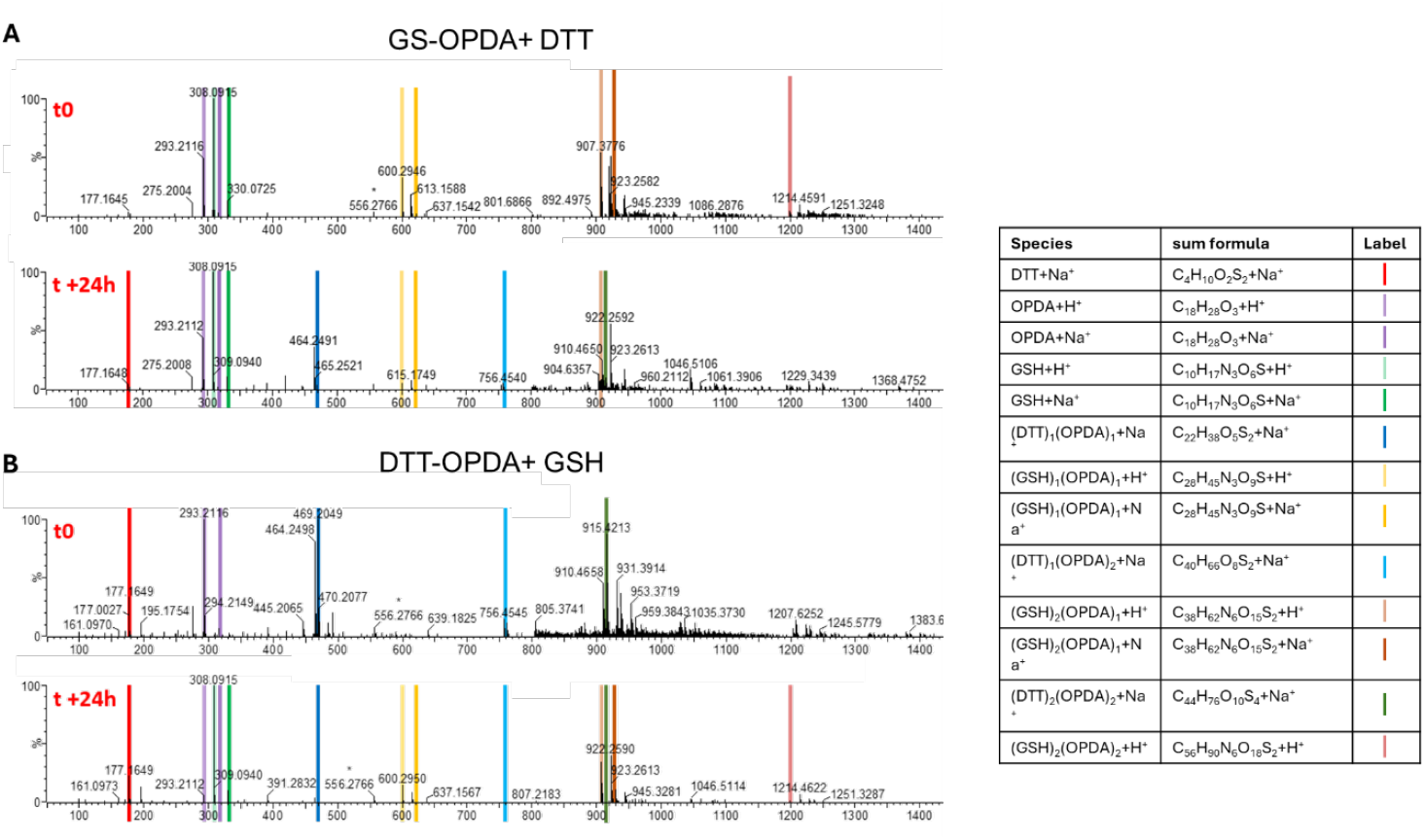
Competition between GSH and DTT for formation of OPDA adducts. 150 µM OPDA was incubated with 1 mM GSH (A) or 1 mM DTT (B) for 16 h at RT. After addition of 1 mM DTT (A) or 1 mM GSH (B) and incubation for up to 24 h (see Tables S3 and S4), formed adducts were assessed by MS. Depicted are measurements after 0 and 24 hours.

To obtain first insights into de-OPDAylation *in vivo*, leaf discs of *A. thaliana* were pre-incubated with 25 µM OPDA, then 1 mM of GSH was added to the floating solution directly before exposure to EEE stress (800 µmol photons m^-2^ s^-1^, 6 h). This test system for OPDA effects on leaf discs had been optimized and characterized, also for its effects on the ascorbate and dehydroascorbate system of the tissue which is the electron donor to ascorbate peroxidases in H_2_O_2_ detoxification (Fig. S2 A). The overnight incubation with OPDA for 16 h and the subsequent EEE treatment for 6 h reliably showed a suppression of the EEE-induced H_2_O_2_ increase with concomitant less oxidation of the ascorbate system (Fig. S2 B). This effect was also seen in the *coi* mutant with defective JA signaling, indicating that this effect is upstream of JA, and, therefore, most likely a direct OPDA effect.

Stress treatment caused a distinct increase in H_2_O_2_ levels, which was absent in leaf discs pretreated with OPDA (Fig. 8A). Addition of GSH to the solution directly before stress exposure partially hindered OPDA-dependent signaling as indicated by a lesser increase in H_2_O_2_. Additionally, measurement of total flavonoid content revealed basal flavonoid levels in mock-treated samples, whereas OPDA pretreatment caused a significant increase (Fig. 8B). In contrast to H_2_O_2_ content, addition of GSH did not lead to an inhibition of this readout of OPDA signaling.

**Figure 8:**
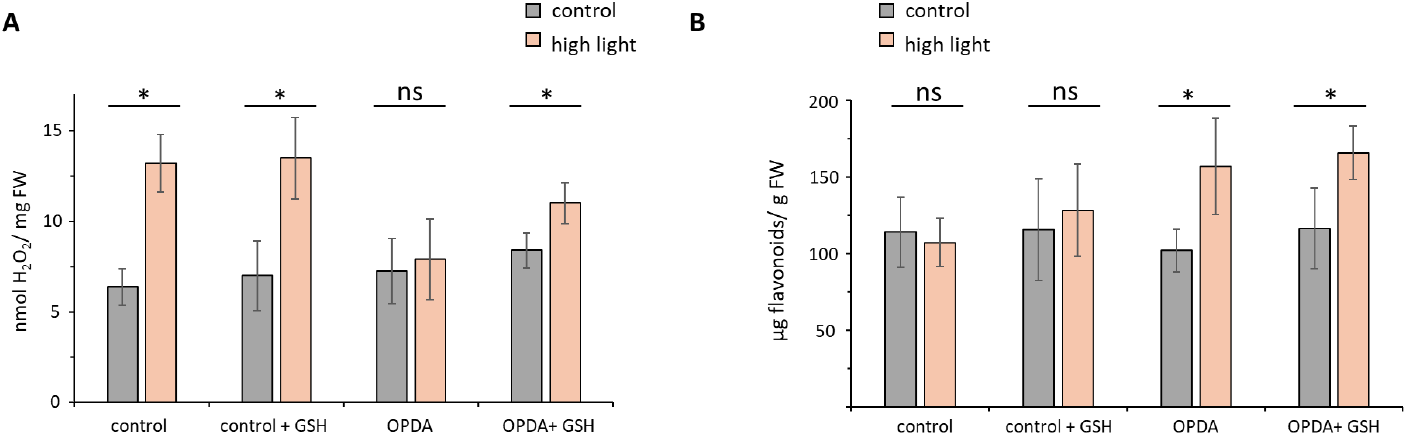
H_2_O_2_ and flavonoid content of leaf discs exposed to high light stress for 6 h. Leaf discs were treated with 25 µM OPDA, 0.27% (v/v) EtOH as control or 1 mM GSH in different combinations to assess the effect of GSH addition on OPDA signaling. Statistically significant increases (p<0.05) in H_2_O_2_ or flavonoid levels after stress treatment were calculated by t test and are denoted as *. Data are means ± SD of n≥5.

Overall, addition of GSH to protein-OPDA adducts regenerated the reduced and active protein form *in vitro*. Occurrence of de-OPDAylation is further supported by GSH-mediated suppression of OPDA-dependent reduction in H_2_O_2_ levels after EEE stress. As de-OPDAylation was quite slow as compared to OPDAylation, enzymatic acceleration *in vivo*, for instance by GSTs, may be hypothesized.

### 3.3 Does GSH function as an efficient competitor for protein OPDAylation?

As GSH majorly influences OPDA signaling by adduct formation as well as de-OPDAylation, a possible antagonistic effect of GSH against OPDAylation of proteins might be assumed. In contrast to the abundant GSH present in mM concentrations, OPDA concentrations remain in the µM range even under stress. If GSH would act as a scavenger of OPDA *in vivo*, GS-OPDA adducts should already form in the chloroplast and any OPDA signaling might be highly unlikely. Hence, our next experiments addressed the question whether OPDAylation of proteins occurs also under physiological concentrations of OPDA (µM) and GSH (mM).

Effects of OPDA on intrinsic fluorescence of TRX-f1 and -m4 was measured in the presence of GSH (Fig. 9, Table 2). As before, OPDA treatment led to a significant decrease in fluorescence intensity (−11.98 and -14.84 for TRX-f1 and m4, respectively), resembling the OPDA effect without GSH (Table 1) and arguing against a protective effect of GSH.

**Table 2:** Intrinsic protein fluorescence of 15 µM TRX-f1 and TRX-m4 incubated with OPDA without or with pretreatment with GSH for 2h. Given values refer to maximal fluorescence intensity (r. U). Data are means ± SD of n=5. Statistically significant differences (p<0.05) of fluorescence intensity after OPDA treatment were calculated by t test and are denoted as *.

| Treatment | TRX-f1 | TRX-m4 |
| --- | --- | --- |
| Control + EtOH | 116.8 ± 1.9 | 115.3 ± 3.1 |
| OPDA | 102.3 ± 1.8 * | 99.0 ± 1.7 * |
| GSH + EtOH | 112.7 ± 0.4 | 116.6 ± 0.7 |
| GSH + OPDA | 100.7 ± 0.7 * | 101.7 ± 0.3 * |

**Figure 9:**
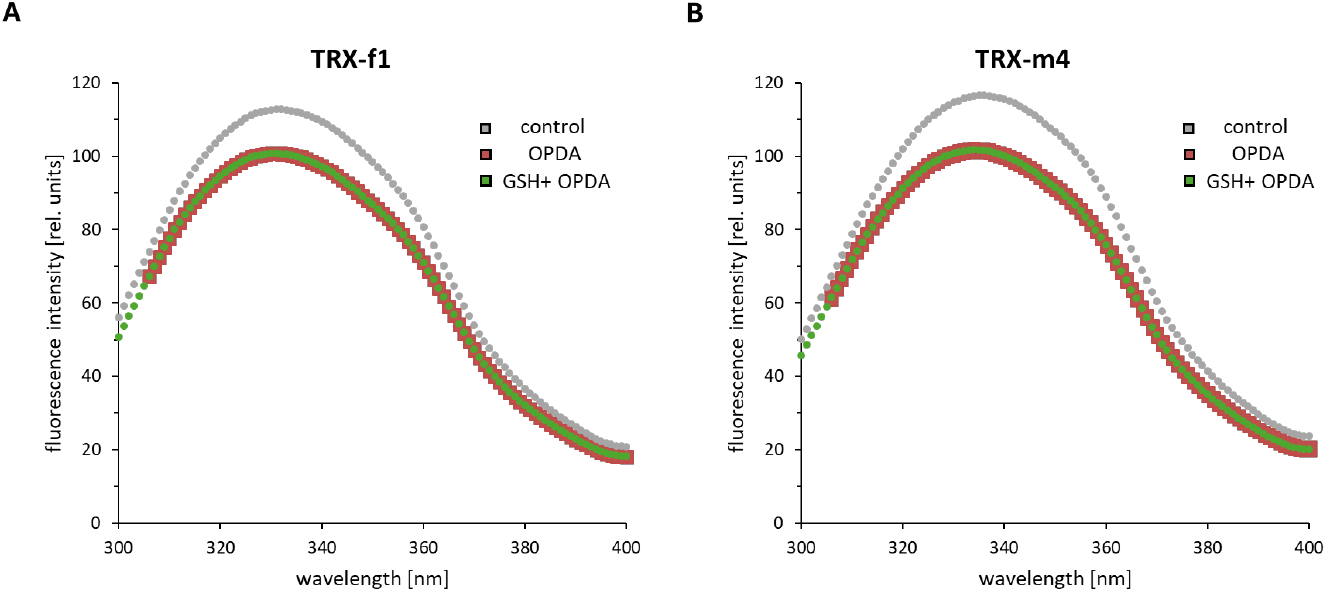
Intrinsic tryptophan fluorescence of TRX-f1 and TRX-m4. Recombinant protein (15 µM) was incubated with 1 mM GSH for 2 h followed by 150 µM OPDA for 3 h. As a control, protein was incubated with GSH and 0.27% (v/v) EtOH. Data are means of n=5.

Additionally, insulin reduction activity of TRX-h3, -f1 and -m4, pretreated with GSH, was analyzed after addition of OPDA (or 0.27% [v/v] EtOH as a control). Again, no effect of GSH on OPDAylation capacity could be observed (Fig. 10). However, after prolonged incubation (16 h), reduction activity of TRX-h3, -f1 and -m4 recovered to 89.7%, 93% and 84.3%, respectively, indicating de-OPDAylation.

**Figure 10:**
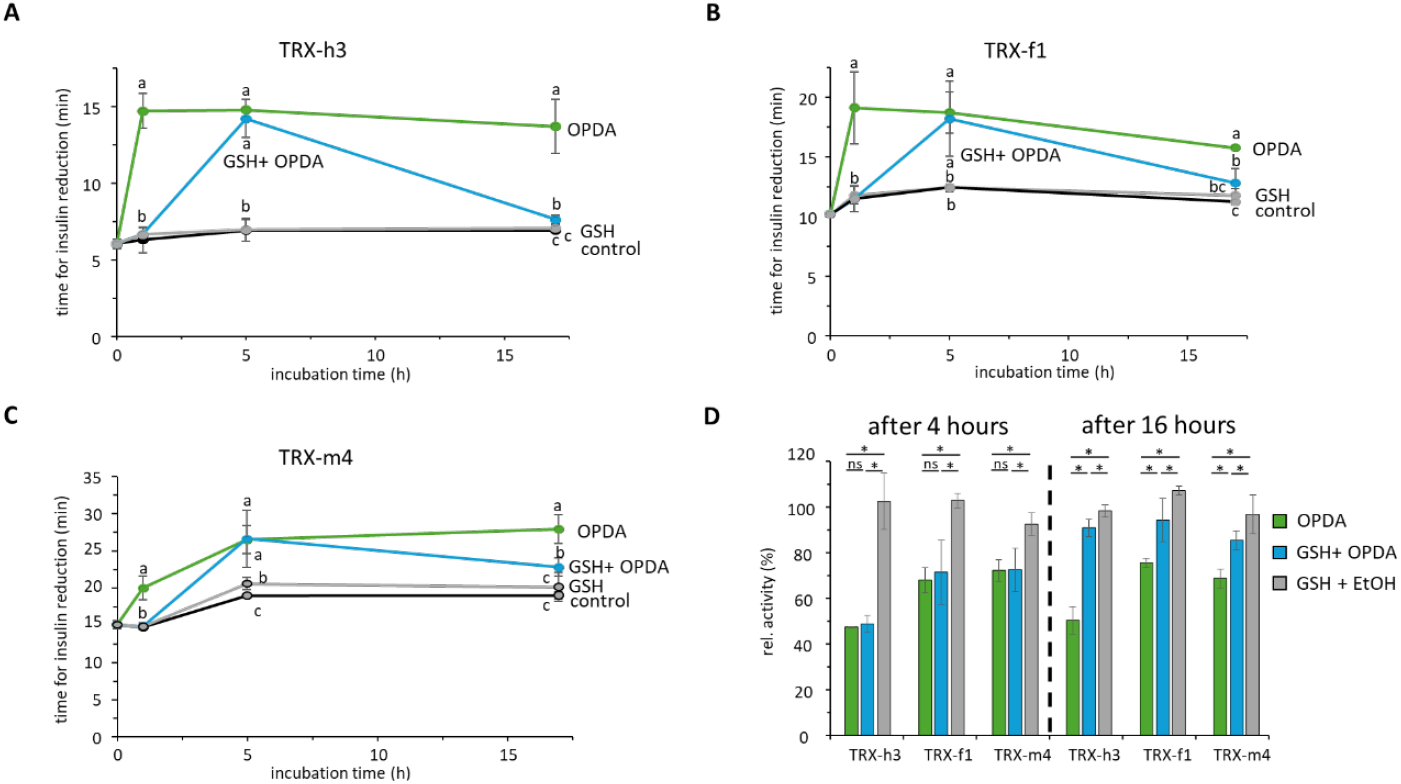
Insulin reduction activity of thioredoxins after pretreatment with GSH. Recombinant TRX (20 µM) was treated with 1 mM GSH for 1 h. Then, 80 µM OPDA or 0.27% [v/v] EtOH were added, and activity was measured after 4 and 16 h. (A-C) Protein activity is depicted as the time needed to pass the threshold of ΔA_650 nm_=0.3. Data are means ± SD of n≥3. (D) Thioredoxin activity was determined as the change in A_650 nm_ min^-1^ using a final protein concentration of 5 µM. Data were normalized to a control sample containing protein incubated only with 0.27% [v/v] EtOH. Significance of difference was assessed by t-test (p<0.05) and denoted either as letters a-c or as *.

Lastly, the effect of GSH pretreatment was analyzed *in vivo* by applying 1 mM GSH to leaf discs and adding 25 µM OPDA after 16 h of incubation. After addition of OPDA, EEE was applied and contents of H_2_O_2_ and flavonoids were measured as described before (Fig. 8). Neither OPDA-dependent changes in H_2_O_2_ nor in flavonoid levels were disturbed by GSH pretreatment (Fig. 11). This further supports the conclusions from the *in vitro* experiments that GSH does not scavenge OPDA before protein-OPDA adducts can form.

**Figure 11:**
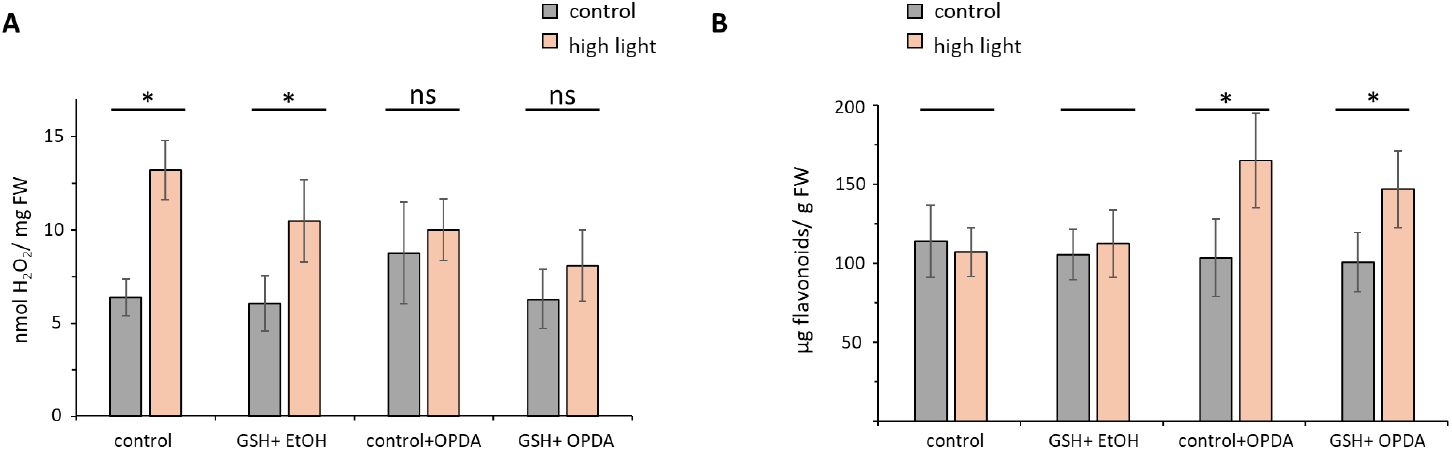
H_2_O_2_ and flavonoid content of leaf discs exposed to EEE stress for 6 h. Leaf discs were pretreated with 1 mM GSH (or H_2_O as a control) for 16 hours. Directly before stress treatment, 25 µM OPDA or (0.27 % [v/v] EtOH) was added to assess the effect of GSH pretreatment on OPDA signaling. Statistically significant increase (p<0.05) in H_2_O_2_ or flavonoid levels after high light were calculated by t-test and are denoted as *. Data are means ± SD of n≥5.

### 3.4 Retro-Michael addition versus thiol exchange mechanism

The data shown so far mark OPDAylation as reversible PTM, with possible release of bound OPDA moieties by excess low molecular weight thiols. However, the exact mechanism underlying de-OPDAylation remains unclear. Hypothetically, adducts could either undergo *retro*-Michael addition liberating OPDA, or OPDA could be transferred directly to another thiol-containing compound by a thiol exchange mechanism that would then follow a nucleophilic substitution mechanism.

To this end, levels of free OPDA were assessed after release of OPDA from TRX-f1 (Fig. 12) and TRX-h3 (Fig. 13). Neither the HPLC-based quantification of OPDA nor the MS analysis showed a return of free OPDA. However, involvement of enzymes dedicated to de-OPDAylation, for instance GSTLs or DHARs, as proposed before, might constitute an additional mechanism of OPDA release by recovering free OPDA.

**Figure 12:**
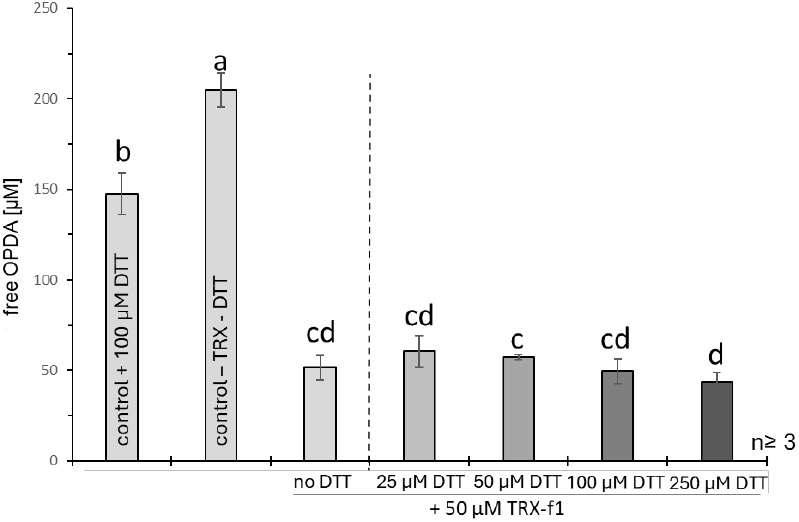
Quantification of OPDA after addition of DTT to OPDAylated TRX-f1. 50 µM TRX-f1 was first incubated with 200 µM OPDA for 2 h, then 0-250 µM DTT was added and OPDA content was analyzed in the sample after 6 h. Controls consisted of TRX-OPDA without DTT, OPDA with only DTT and OPDA without additional compounds. Data are means ± SD of n=3, significant differences were assessed by one-way ANOVA and Tukey and are marked as a-d.

**Figure 13:**
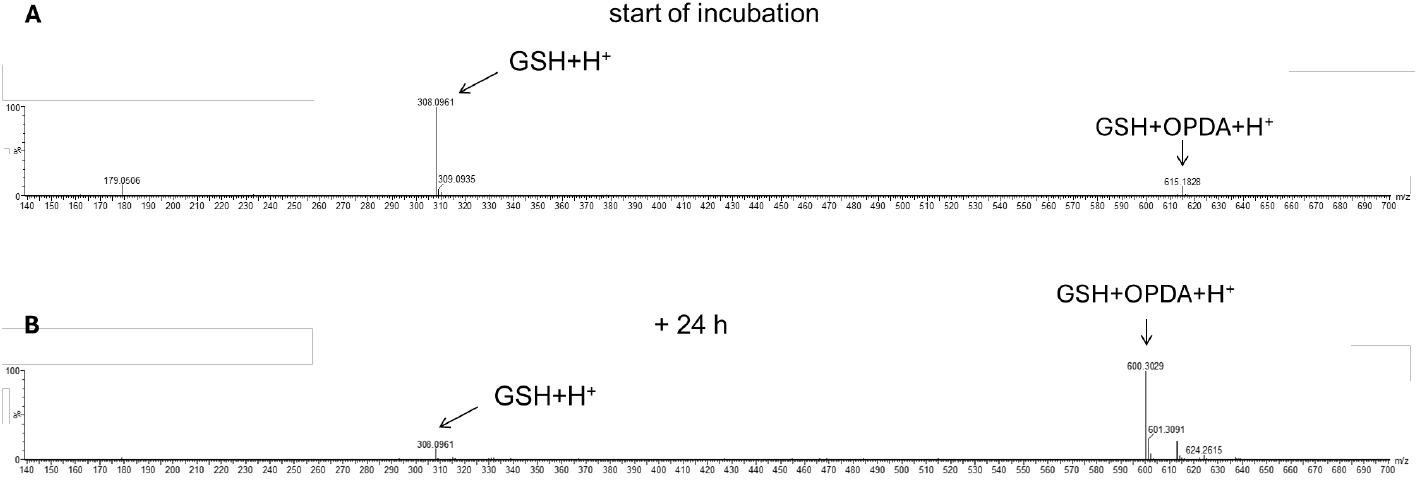
MS analysis of free OPDA, GSH and GS-OPDA. MS analysis (corresponding to Fig. 3) was performed after addition of GSH to desalted TRX-h3-OPDA adducts over a time span of 24 h. GS-OPDA adduct was detectable from 4 h onward. After 24 h, free GSH+H^+^ and GS-OPDA, but no OPDA could be detected.

## 4. Discussion

OPDAylation has been proposed as a novel posttranslational modification regulating the activity of thiol-containing proteins of the redox regulatory network [10]. Through this mechanism, OPDA could influence redox homeostasis and subsequently gene expression *in planta* to promote stress defense. This study aimed at clarifying whether OPDAylation is a stable or reversible PTM by combining a variety of *in vitro* and *in vivo* studies.

### OPDAylation as a reversible process

The results of this study prove that OPDAylation of proteins is reverted in the presence of excess low molecular mass thiol compounds (GSH or DTT). The results show de-OPDAylation for different proteins with established roles in the redox regulatory network, including the cytosolic TRX-h3, plastid-localized TRX-f1 and -m4, and Cyp20-3. De-OPDAylation recovered the protein-specific activity, namely the insulin reduction ability and electron donor capacity to GPXL8 as indicated by the H_2_O_2_ reduction (Figs. 4, 5). In general, the extent of recovery increased in proportion to the protein’s sensitivity to OPDAylation-linked inhibition. For instance, PPIase activity of Cyp20-3, which is highly sensitive towards OPDAylation, recovered by 154.1% after 4 h (Fig. 6C, at pH 7.2), while TRX-m4 activity only increased by 12.6% (Fig. 4). Interestingly, like spontaneous GS-OPDA formation, de-OPDAylation was pH-dependent with maxima at pH 6.2 and pH 8.2 (227% and 256%, respectively, Fig. 6C). This might contribute to spatial regulation of de-OPDAylation as adducts could be more stable in cell organelles with neutral pH (ER, cytosol, nucleus) compared to slightly alkaline compartments represented by the mitochondrial matrix, peroxisomes and the chloroplast stroma under illumination [51].

These findings are in accordance with a first-order-rate elimination from a conjugate base (E1cB mechanism), which has been reported for elimination processes of thioether products obtained from Michael addition reactions with thiols such as GSH and its replacement with methyl mercaptane [52].The rate determining step in such an elimination is the formation of the enolate anion, and this step is favored under more basic conditions. Accordingly, a higher degree of de-OPDAylation can be expected at an increased pH value, which is in agreement with the experimental observations described above. In analogy to the E1cB mechanism reported in literature for conversion of glutathione-substituted Michael adducts to MeS-substituted analogues [52], a proposed reaction mechanism is given in Scheme 1 for the de-OPDAylation of the OPDA-adduct with TRX and OPDA formation under release of the thiolate anion of TRX.

**Scheme 1:**
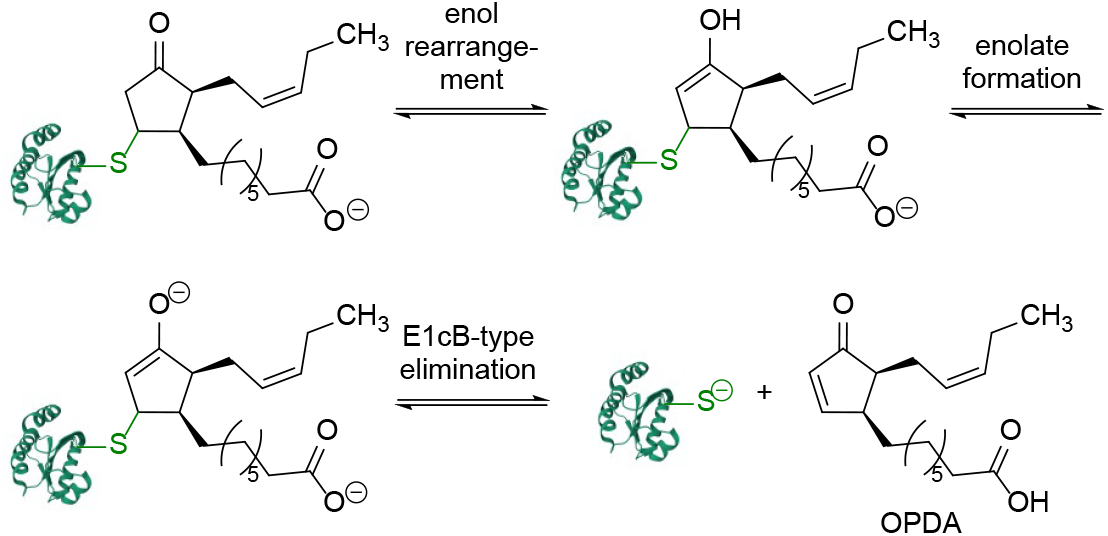
Proposed mechanism for the de-OPDAylation of an OPDA-adduct of thioredoxin (Trx). Cleavage of the OPDA-adduct of thioredoxin proceeding through the steps of enol rearrangement, deprotonation and subsequent E1cB-type elimination; PDB reference of the thioredoxin (from *A. thaliana*; reduced form): 1XFL.

### Protein OPDAylation is favored over OPDA adduct formation with GSH

Under physiological conditions, the concentration of free thiols, for example GSH, exceeds that of OPDA about 10^2^-to 10^3^-fold. The fact that these thiols react with OPDA raises the question how basal levels of free OPDA are maintained for OPDA signaling as well as for processing to jasmonic acid. Remarkably, OPDAylation of TRX-f1 and TRX-m4 occurred *in vitro* independent of the presence of GSH, as indicated by OPDA-dependent quenching of intrinsic Trp fluorescence and decreased insulin reduction activity (Table 1, 2, Fig. 10). *In vivo* studies on OPDA signaling during EEE further supported the occurrence of OPDAylation under physiological conditions (Fig. 11).

As depicted in Fig. 2, spontaneous GS-OPDA formation is a slow process. For instance, after 4 h of incubation of 250 µM GSH and 500 µM OPDA at pH 7.2, ∼50% of GSH was bound to OPDA. Further ^1^H NMR spectroscopic studies with an excess of GSH (ratio of GSH to OPDA of 5:1) supports these findings of a relatively slow OPDAylation step with GSH as the thiol component as indicated by a conversion of 59% after 3 h when starting from a concentration of OPDA of 5 mM. In contrast, protein OPDAylation proceeds rapidly, modifying activity of target proteins distinctly already after 10 min [12, 10]. Different degrees of OPDA-mediated inhibition of distinct proteins indicate specificity. Both factors appear to contribute to preferential binding of OPDA to protein thiols instead of GSH. However, various GSTs catalyze GS-OPDA binding *in vivo* and will accelerate the reaction as discussed below [10, 11, [53] [54, 34].

These results can be rationalized by the hypothesis that initial TRX-based OPDAylation is proceeding under kinetic control due to the higher reactivity of TRX compared to GSH. The higher nucleophilicity of proper TRX is due to the cysteine moiety C32 being present in TRX as a part of the active site of the reduced form of TRX, namely the Cys32-Gly-Pro-Cys35 motif [55].This molecular environment facilitates deprotonation of C32, which then reacts as a thiolate [55],thus leading to a much faster kinetically controlled Michael addition compared to the thiol moiety in GSH. However, due to the excess of GSH being present in the cell at much higher concentrations compared to TRX, at prolonged reaction time the OPDAylation of GSH is observed as a result of a thermodynamically controlled process. The relationship of these two types of reversible OPDAylation reactions with the sulfur nucleophiles TRX and GSH, respectively, is visualized in Scheme 2.

**Scheme 2:**
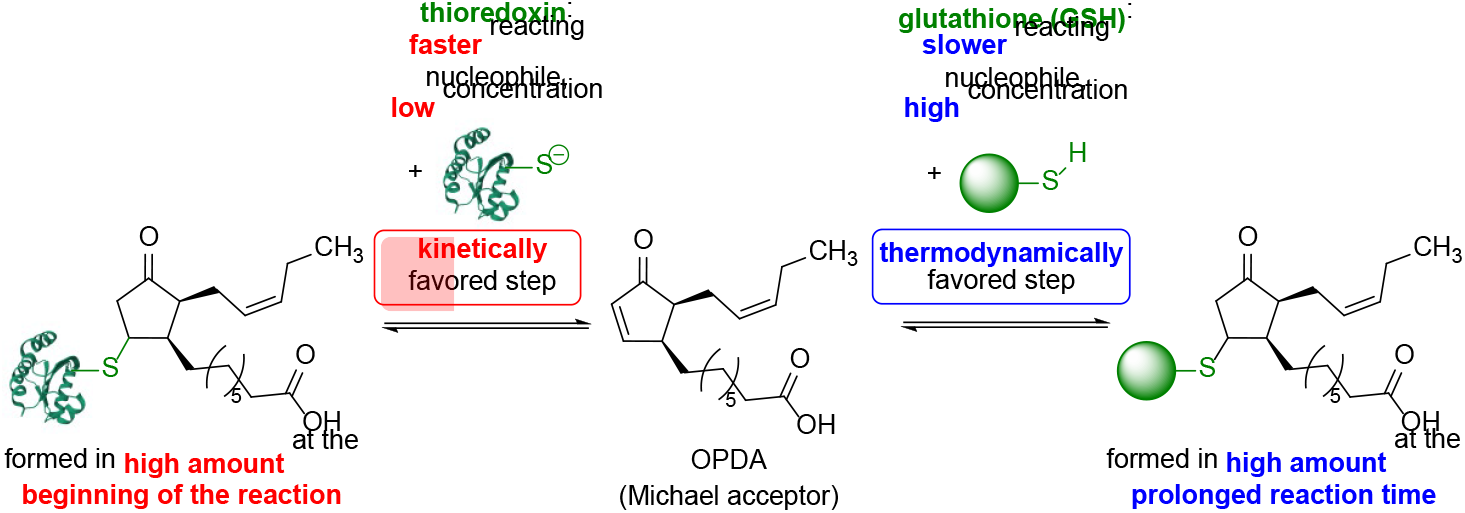
Kinetically versus thermodynamically controlled reaction for the formation of *OPDA*-adducts of thioredoxin (TRX) and glutathione (GSH), respectively. Initially the preferred formation of the OPDA-adduct of TRX prevails due to the fast, kinetically controlled addition of the thiolate in TRX and thermodynamically controlled formation of the OPDA-adduct of GSH as preferred adduct at prolonged reaction time due to the large excess of GSH; PDB reference of the TRX (from Arabidopsis thaliana; reduced form): 1XFL.

### OPDA signaling under high light stress

The initial predominance of the OPDAylation process of proteins in the presence of GSH, followed by (slow) de-OPDAylation, as observed *in vitro*, revealed novel dynamics of OPDA signaling. To scrutinize these findings *in vivo*, leaf discs pretreated with OPDA and/or GSH were subjected to EEE for 6 h followed by determination of H_2_O_2_ and flavonoid content. OPDA positively regulates both the degree of ROS production and the cellular ROS detoxification capacity by stimulating the synthesis of antioxidants, chaperones and photoprotective pigments in a Cyp20-3 dependent manner [17, 11, 56, 22, 23, 21].

Publicly available transcriptome data of *A. thaliana* subjected to EEE, heat stress or wounding (Fig. S3) reflects general upregulation of GSTs (and jasmonate synthesis) under non-optimal growth conditions. However, affected GSTs seem to vary between stress type (and duration), creating specific GST signatures. For instance, GSTL1, GSTU2 and GSTU10 are more strongly upregulated after wounding, whereas DHAR3 and GSTU20 rather respond to high light stress together with MYB transcription factors MYB11 and MYB111. GSTU20, an identified target of OPDA signaling [53], as well as MYB11/MYB111 positively influence flavonoid synthesis and stability [57–59].

Leaf discs pretreated with 25 µM OPDA lacked the EEE-induced increase in H_2_O_2_ observed in control samples. Instead, total flavonoid content was significantly enhanced, arguing for OPDA-mediated acceleration of flavonoid biosynthesis (Fig. 11). Both readouts remained stable when leaf discs were pretreated first with GSH and secondly with OPDA. This result supports the preferential interaction of OPDA with protein thiols, specifically Cyp20-3, previously observed *in vitro*.

However, de-OPDAylation, as tested by addition of GSH after OPDA-pretreatment, recovered only the EEE-induced H_2_O_2_ accumulation, while flavonoid accumulation was not affected (Fig. 8). Although this appears contradictory at first, adjustment of flavonoid levels is much less dynamic compared to ROS levels, with first measurable increases after 12 to 24 h or 10 to 60 min, respectively [58, 60, 19, 61]. Hence, modulation of H_2_O_2_ content can be regarded as an indicator of maintenance of OPDA signaling, while flavonoid levels display OPDA signaling in the early stage of EEE. Consequently, it might be assumed that excess external GSH did not impede initial OPDAylation of target proteins and resulting ORG expression, but duration of OPDA signaling was shortened.

A possible reason might be a disturbance in the GS-OPDA-to-OPDA ratio. GSTs accumulate already in early EEE stress, [19]. Additionally, the stromal pH increases from pH 7 to more than 8 under illumination [62]. As both GSH binding and de-OPDAylation showed a clear pH dependency, this might increase the GS-OPDA-to-OPDA ratio in EEE further. Formation of GS-OPDA (and detoxification of ROS) is GSH-consuming, resulting in a decreased GSH-to-GSSG ratio. This decrease, and the simultaneous increase in GS-OPDA, are measurable after 15 min of EEE stress [19]. As gene expression of MRP2 (EC:7.6.2.2) and GGT4 is not yet upregulated (Fig. S3), GS-OPDA accumulation most likely serves as a means of OPDA transport and/or JA synthesis, and not degradation. Although JA content under EEE is not significantly enhanced in the long-term [21], controlled conversion of OPDA to JA would allow for JA-dependent stimulation of jasmonate synthesis. This would be in line with enhanced gene expression of OPR3 and OPCL1 (EC:6.2.1.-) (Fig. S3) and minor increases in JA content, which are observable after 15 to 60 min [19]. The pool of reduced GSH available for GS-OPDA formation might be a limiting factor at this point, preventing complete depletion of free OPDA. By supplementing GSH in the floating solution, however, this bottleneck might be avoided, resulting in increased turnover of OPDA and subsequent loss of OPDA signaling.

### Changes in cellular thiol composition modulate OPDA signaling

Based on the interplay of protein OPDA adducts on the one hand and GS-OPDA on the other hand, the following hypothetical scheme of OPDA signaling dependent on variations in cellular thiol composition might be deduced (Fig. 14).

**Figure 14:**
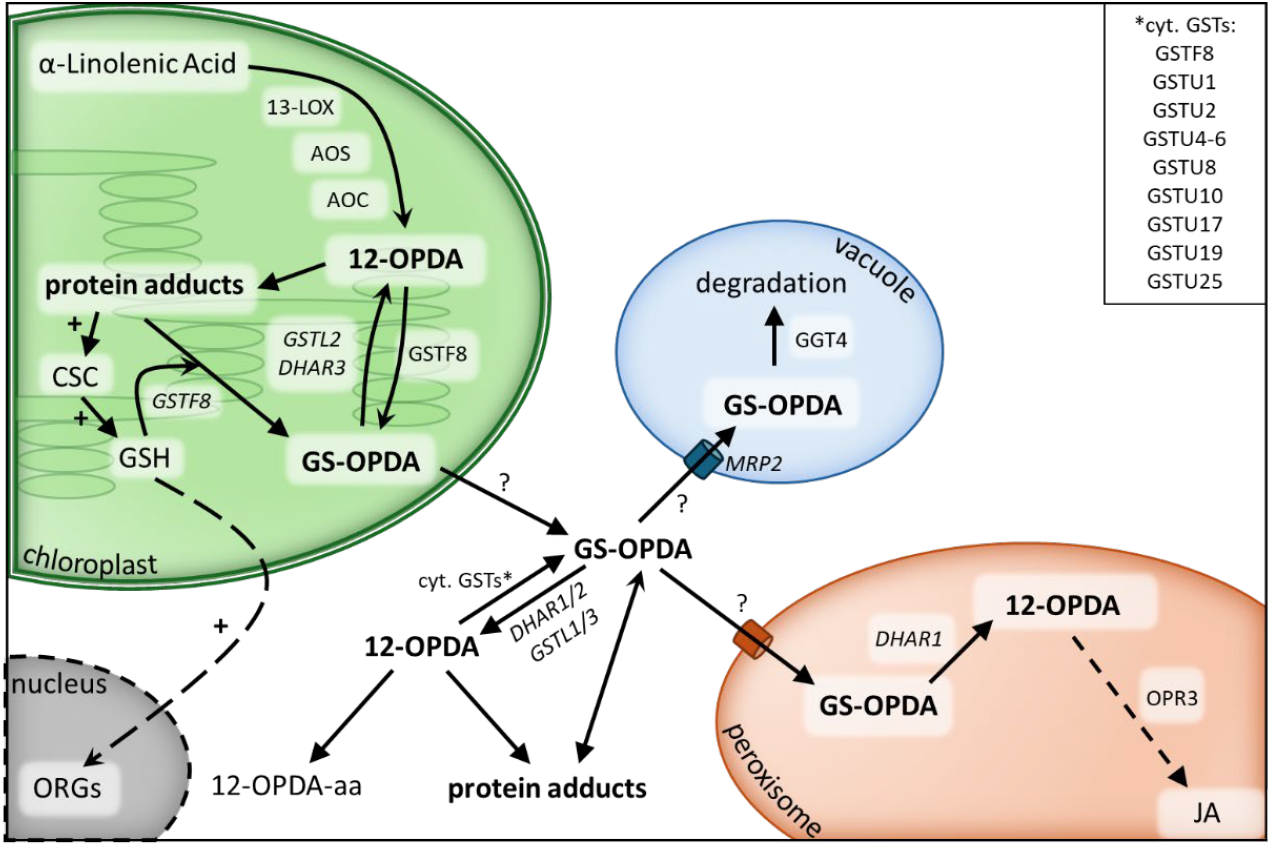
Availability of OPDA for signaling, degradation or conversion to JA might be tightly linked to interaction with GSH. OPDA generated in the chloroplast forms protein-OPDA or GS-OPDA adducts. While transfer of OPDA moieties between adducts is possible, free OPDA might be regenerated by DHARs and GSTLs. GS-OPDA formation, transport and regeneration of OPDA depends on expression pattern of GSTs. As interaction with Cyp20-3 stimulates GSH synthesis and ORG (OPDA-regulated genes) expression, GST-mediated OPDA modulation likely is a negative feedback system. Enzymes written in italics (GSTLs/DHARs) have not been analyzed for activity on GS-OPDA so far.

Upon synthesis in the chloroplast OPDA likely binds to proteins such as Cyp20-3. The alkaline pH of the stroma established in the light may foster this reaction [63]. The adduct formation initiates OPDA signaling via activating the Cysteine synthase complex and enhanced O-acetylserine, Cys and subsequently glutathione synthesis [64]. Considering the dynamic interplay between OPDAylation and de-OPDAylation and the role of free thiols, the accumulation of GSH may be considered as feedback inhibitory circuitry ending or decreasing the OPDA-dependent signaling in the chloroplast.

Free protein recovers by transfer of bound OPDA moieties to GSH, either spontaneously or, possibly, catalyzed by GSTs. While this transfer might occur in both directions, there is no regeneration of free OPDA in the spontaneous reaction (Figs. 12, 13). However, unbound OPDA is crucial for JA synthesis, as GS-OPDA cannot serve as substrate for OPR3 [43]. It may be hypothesized that GSTs of the DHAR and Lambda class, which can act as thioltransferases and show activity towards GS adducts [65, 38] catalyze this reaction. Like other GSTs, DHAR1 expression is regulated by OPDA and additionally by JA [17, 65].

Based on the subcellular localization of OPDA-active GSTs in the cytosol and plastid stroma and GSTLs and DHARs in plastids, cytosol, peroxisome and mitochondria, GST-dependent modulation of the GSH-OPDA interaction could serve as a means of OPDA transport. A similar mechanism of GSH-mediated transport has already been described for animal prostaglandins [36, 66, 67]. Along this line, plastid OPDA could be bound to GSH by GSTF8 and transported into the cytosol (e.g. through ABC transporters) where it could target cytosolic proteins or be bound to amino acids after GSTL- or DHAR-mediated adduct dissociation.

This hypothetical mechanism might control OPDA detoxification as a negative feedback system, as OPDA signaling causes upregulation of GST gene expression. With increasing GST levels, the GS-OPDA-to-OPDA ratio likely increases as well. GS-OPDA could then be transported either to the vacuole for degradation or to the peroxisome for OPDA release by DHAR1 and subsequent JA synthesis. Thereby, the dynamic interplay of OPDA with protein thiols and GSH might fine-tune OPDA-signaling, degradation and metabolisation.

## 5. Conclusions

OPDAylation targets essential components of the plant redox regulatory network with severe functional implications. This observation motivated us to conduct this research with two main aims: First, OPDAylation was further scrutinized, and the results prove OPDAylation as reversible PTM. De-OPDAylation depends on the protein’s sensitivity towards OPDA, the pH and the concentration of excess thiol groups. This likely allows for precise modulation of OPDA amounts, as the rapid modification of protein activity enables subsequent induction of OPDA signaling, whereas de-OPDAylation, triggered by increasing GSH, increasing cellular reduction potential (more negative redox potential) or enzymatically by GSTs, reduces oxidative damage which might be caused by pronounced alteration of the thiol switch proteins. It will be important in the future to explore the products of GS-OPDA cleavage.

The involvement of GSH in protein recovery prompted the second focus of this work. Owed to its high abundance, GSH could act as an OPDA scavenger, modulating or preventing OPDA signaling and JA synthesis. However, OPDA signaling could be detected even in the presence of GSH. This interplay of OPDAylation and de-OPDAylation might indicate a GSH-dependent network regulating OPDA modification and signaling. GSH-dependent modulation of OPDA signaling might contribute to the plant response to different stresses, especially those for which uncoupling of OPDA-from JA-signaling has been proposed [68, 19, 69, 3]. However, plant stress defense is highly complex and varies not only based on the type of stress, but also on other factors in particular stress duration, stress intensity, developmental state of the plant and on the combination of simultaneously occurring stresses [70]. Future studies should explore the involvement of DHARs and GSTLs in OPDAylation and de-OPDAylation.

## Abbreviations

2-CysPrx: 2-Cysteine peroxiredoxin
4-HNE: 4-Hydroxynonenal
ABC: transporter ATP-binding cassette transporter
ANOVA: Analysis of variance
AOC: Allene oxide cyclase
AOS: Allene oxide synthase
CSC: Cysteine synthase complex
Cyp20-3: Cyclophilin 20-3
DHAR: Dehydroascorbate reductase
dn-OPDA: Dinor-OPDA
DTNB: 5,5’-Dithiobis-2-nitrobenzoic acid
DTT: Dithiothreitol
EEE: Excess excitation energy
GEO: Gene Expression Omnibus
GGT: γ-Glutamyl transpeptidase
GPXL: Glutathione peroxidase-like
GSH: Glutathione
GST: Glutathione-*S*-transferase
JA: Jasmonic acid
KPi: Potassium phosphate
LOX: Lipoxygenase
MRP: Multidrug resistance-associated protein
MS: Mass spectrometry
MYB: Myeloblastosis
NCBI: National Center for Biotechnology Information
NTRA: NADPH-dependent thioredoxin reductase A
OPC:8-0: 8-(3-Oxo-2-(pent-2-enyl)cyclopentenyl)octanoid acid
OPDA: *Cis*-(+)-12-oxophytodienoic acid
OPR: OPDA reductase
ORG: OPDA-responsive gene
PPIase: eptidyl-prolyl-*cis*-*trans*-isomerase
PTM: Posttranslational modification
r.u.: Relative units
RCS: Reactive carbonyl species
ROS: Reactive oxygen species
TRX: Thioredoxin
WT: Col-0 Wild-type Columbia-0
α-LeA: α-Linolenic acid

## Author contributions

Conzeptualization, A.V., M.K., K.-J.D.; methodology and investigation, A.K., J.S., L.V., M.K., R.S.; resources, H.G., M.B.; writing-original draft preparation, M.K.; review and editing, all authors; supervision, K.-J.D.; funding acquisition, H.G., K.-J.D. All authors have read and agreed to the published version of this manuscript.

## Acknowledgements

We thank Tim Guntelmann (Industrial Organic Chemistry and Biotechnology, Bielefeld University, Germany) for providing OPDA in the initial phase of the project.

## Supporting information

Table S1: MS analysis of TRX-h3-OPDA adducts using different OPDA:protein ratios.

Table S2: Stability of TRX-h3:OPDA adducts over 24 hours as determined by MS.

Figure S1: ^1^H-NMR analysis of GS-OPDA adducts.

Figure S2: MS^2^ of GS-OPDA adducts.

Figure S3: MS analysis: GSH-triggered de-OPDAylation of TRX-h3.

Figure S4: Effect of OPDA on EEE-induced H_2_O_2_ accumulation and ratio of ascorbate (AsA) to total ascorbate pool (AsA + dehydroascorbate (DHA).

Figure S5: Expression profiles of selected genes after stress treatment.

