## Supplemental Materials for "The competitive interplay of 12-oxophytodienoic acid (OPDA), protein thiols and glutathione"

### Supplementary methods

*Plant growth*

*A. thaliana* Col-0 and *coi1* (kindly provided by Dr. Roberto Solano) plants were grown in soil in a climate chamber with a day/night cycle of 10h/14h with 80 µmol photons $m^{-2}$ $s^{-1}$, 21°C/14°C and 50% relative humidity. Prior to that, seeds were incubated at 4°C for 3 d. After 6 to 7 weeks of growth, leaf discs of 5 mm diameter were cut out using a cork borer and immediately placed upside down in a 96-well-plate filled with either 25 µM oxylipin solution (containing 0.1 mM $CaCl_2$) or, as a mock control, 0.083% [v/v] EtOH solution (also containing 0.1 mM $CaCl_2$). After 16 h of incubation, leaf discs were subjected to EEE stress (800 µmol photons $m^{-2}$ $s^{-1}$) for 6 h and frozen in liquid nitrogen for determination of hydrogen peroxide (see 2.12) and (dehydro-) ascorbate content. Frozen leaf discs were used for analysis within approximately 3-4 days.

*Measurement of ascorbate and dehydroascorbate*

The ascorbate content of leaf discs was determined using a plate reader according to method used in [1] with minor modifications. 10 mg of leaf discs were extracted in 100 µL of HCl (0.2 N), and the extract was neutralized with 0.2 M NaOH. The sample mixture composed of 45 µL of the neutralized extract, 57.9 µL of 0. 12 M sodium phosphate buffer (pH 7.5) and 4.3 µL of 25 mM DTT (for total ascorbate) or $H_2O$ (for reduced ascorbate). Total and reduced ascorbate contents were determined based on the decrease in absorbance at 265 nm using of 5 µL of ascorbate oxidase (0.05 U/µl). The amount of dehydroascorbate was determined as the difference between total ascorbate and reduced ascorbate. The calculations were performed using a corrected extinction coefficient of 7,000 $M^{-1}$ $cm^{-1}$.

*Transcriptome analysis*

To analyze differential expression of selected genes under high light, heat or wounding stress, RNAseq data was obtained from the NCBI GEO repository and iDEP.96 was used for identification of differential gene expression [2]. Data sets used were GSE277977 [3], GSE101422 [4] and GSE134391 [5]. Additionally, transcriptome data generated by Crisp et al. (2017) and Xu et al. (2024) was included [6, 7].

*H-NMR spectroscopic monitoring of GS-OPDA-formation*

To monitor the formation of GS-OPDA, 1H-NMR-spectroscopic measurements were performed on a Bruker Avance III 500 and a Bruker Avance III 500HD. The solvent signal used for the internal calibration of the NMR spectra was D2O (δ = 4.79) [8]. For the NMR-experiment, 12-OPDA (1.2 mg, 0.004 mmol, 1 eq.) and GSH (6.3 mg, 0.020 mmol, 5 eq.) were dissolved in D2O (800 µL). In order to increase the solubility of 12 OPDA, the pH of the solvent was adjusted to pH 8 using sodium hydroxide similar to the stromal pH in illuminated chloroplasts. The sample was incubated at RT and measured at 10, 50, 140, 170 and 180 minutes after preparation.

**Table S1: MS analysis of TRX-h3-OPDA adducts using different OPDA:protein ratios.** 20 µM of recombinant protein was incubated with 0-80 µM of OPDA at RT for 16 hours.

| OPDA:TRX ratio | Base peak | Peak 1 | Peak 2 | Peak 3 |
|---|---|---|---|---|
| 0:1 | 15.14 ± 0.04 | - | - | - |
| 1:1 | 15.14 ± 0.03 | 15.43 ± 0.03 | 15.73 ± 0.2 | - |
| 2:1 | 15.14 ± 0.03 | 15.43 ± 0.02 | 15.73 ± 0.05 | - |
| 3:1 | 15.14 ± 0.03 | 15.43 ± 0.05 | 15.73 ± 0.06 | - |
| 4:1 | 15.14 ± 0.06 | 15.43 ± 0.06 | 15.73 ± 0.08 | 16.02 ± 0.28 |

**Table S2: Stability of TRX-h3:OPDA adducts over 24 h as determined by MS.** Base peak equals free TRX, peak 1-3 resemble 1-3 moieties of OPDA bound to TRX.

| OPDA: TRX | base peak | peak 1 | peak 2 | peak 3 |
|---|---|---|---|---|
| t0 | 15.14 ± 0.06 *10$^{-3}$ kDa | 15.43 ± 0.02 *10$^{-3}$ kDa | 15.73 ± 0.05 *10$^{-3}$ kDa | - |
| t3 | 15.14 ± 0.03 *10$^{-3}$ kDa | 15.43 ± 0.04 *10$^{-3}$ kDa | 15.73 ± 0.07 *10$^{-3}$ kDa | - |
| t6 | 15.14 ± 0.07 *10$^{-3}$ kDa | 15.43 ± 0.02 *10$^{-3}$ kDa | 15.73 ± 0.1 *10$^{-3}$ kDa | 16.02 ± 4.66 *10$^{-3}$ kDa |
| t24 | 15.14 ± 0.05 *10$^{-3}$ kDa | 15.43 ± 0.03 *10$^{-3}$ kDa | 15.73 ± 0.02 *10$^{-3}$ kDa | 16.02 ± 5.15 *10$^{-3}$ kDa |

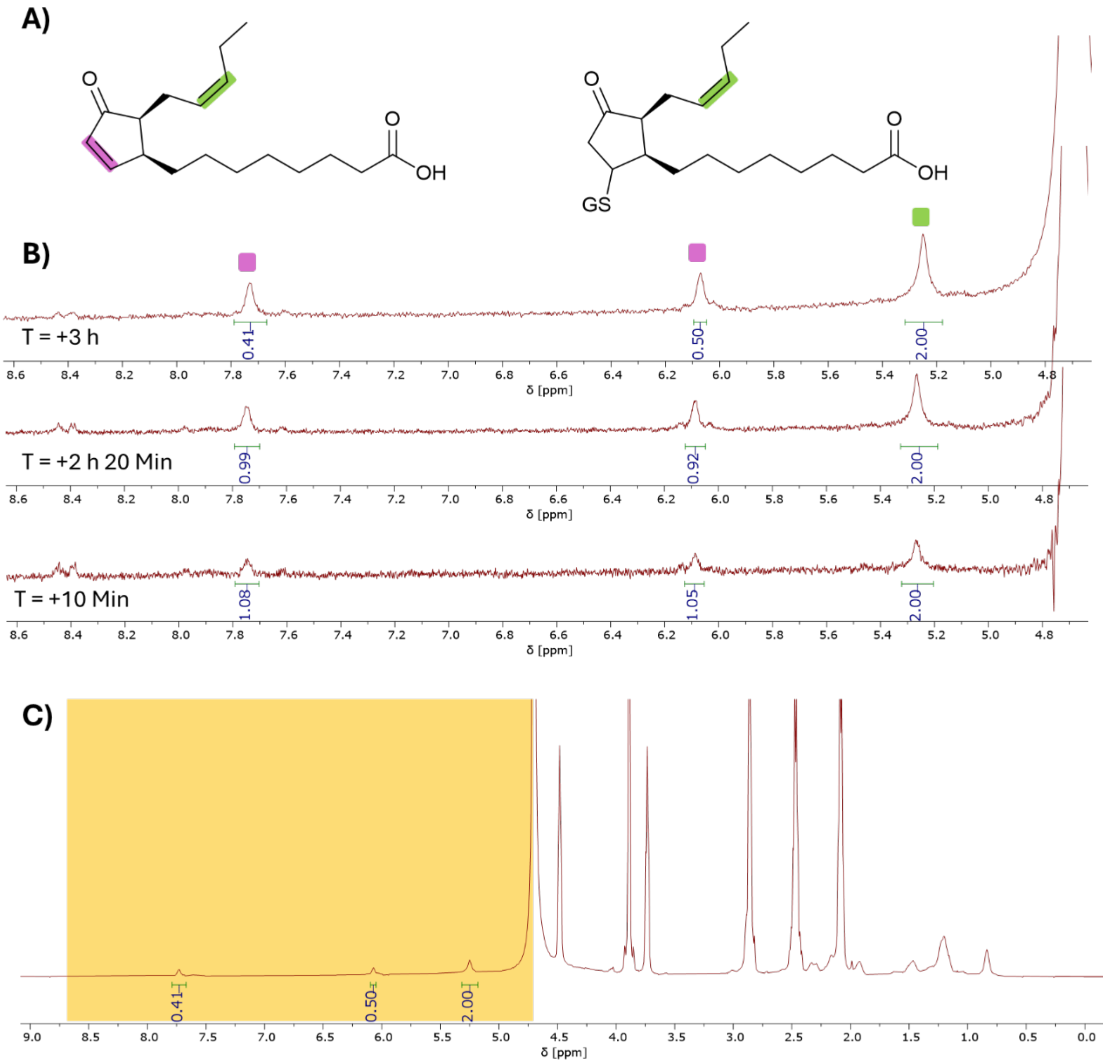


**Figure S1: A)** Structures of 12-OPDA and GS-OPDA. The double-bond protons of the cyclopentene ring and the α-side chain have been marked in pink and green, respectively. **B)** Comparison of $^{1}$H-NMR spectra taken at different time points. The signals are assigned by colour. The formation of the

12-OPDA-glutathione-adduct is indicated by the disappearance of the signals of the cyclopentene-double bond. **C)** Full $^{1}$H-NMR-spectrum recorded at timepoint T = + 3 h. The zoomed-in section from B) is highlighted in yellow.

**A)**

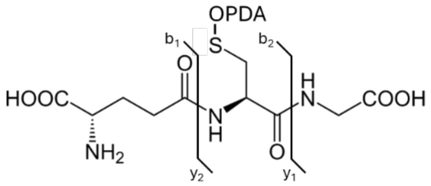


**B)**

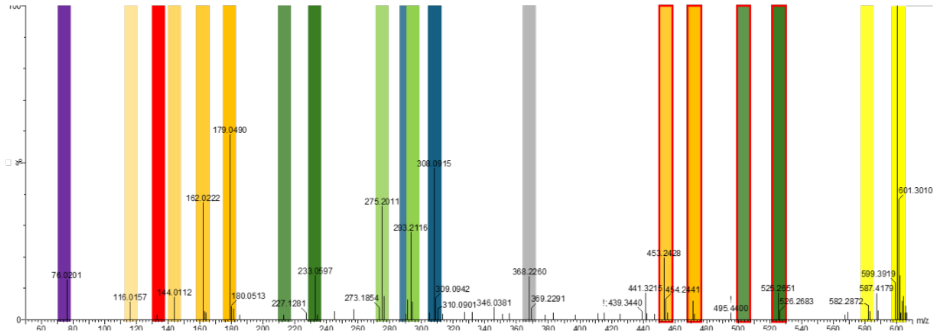


**C)**

| Species | sum formula | $m/z_{(theo)}$ | $m/z_{(exp)}$ | Δ [mmu] |
|---|---|---|---|---|
| GSH+OPDA+H⁺ [M+H⁺] | $C_{28}H_{45}N_3O_9S+H^+$ | 600.2950 | 600.2979 | 2.97 |
| (M-$H_2O$+H)⁺ | $C_{28}H_{43}N_3O_8S+H^+$ | 582.2844 | 582.2872 | 2.87 |
| $b_2$ | $C_{26}H_{40}N_2O_7S+H^+$ | 525.2629 | 525.2651 | 2.20 |
| $b_2$-$H_2O$ | $C_{28}H_{43}N_3O_8S+H^+$ | 507.2523 | 507.2536 | 1.27 |
| $b_2$-OPDA | $C_8H_{12}N_2O_4S+H^+$ | 233.0591 | 233.0597 | 0.65 |
| $b_2$-$H_2O$-OPDA | $C_8H_{10}N_2O_3S+H^+$ | 215.0485 | 215.0489 | 0.41 |
| $y_2$ | $C_{23}H_{38}N_2O_6S+H^+$ | 471.2523 | 471.2538 | 1.47 |
| $y_2$-$H_2O$ | $C_{23}H_{36}N_2O_5S+H^+$ | 453.2418 | 453.2428 | 1.03 |
| $y_2$-OPDA | $C_5H_{10}N_2O_3S+H^+$ | 179.0485 | 179.0490 | 0.51 |
| $y_2$-$NH_3$-OPDA | $C_5H_7NO_3S+H^+$ | 162.0219 | 162.0222 | 0.26 |
| $y_2$-$NH_3$-$H_2O$-OPDA | $C_5H_5NO_2S+H^+$ | 144.0114 | 144.0112 | 0.62 |
| $y_2$-$NH_3$-$H_2O$-CO-OPDA | $C_4H_5NOS+H^+$ | 116.0165 | 116.0157 | 0.76 |
| $b_1$ | $C_5H_7NO_3+H^+$ | 130.0499 | 130.0512 | 1.33 |
| $y_1$ / Cys-Imin-Fragment | $C_2H_5NO_2+H^+$ <br> $C_2H_5NS+H^+$ | 76.0393 <br> 76.0216 | 76.0395 <br> 76.0201 | 0.20 <br> 1.45 |
| Cys-Imin-Fragment + OPDA | $C_{20}H_{33}NO_3S+H^+$ | 386.2254 | 368.2260 | 0.61 |
| M-OPDA+H⁺ = GSH+H⁺ | $C_{10}H_{17}N_3O_6S+H^+$ | 308.0911 | 308.0915 | 0.42 |
| M-OPDA-$NH_3$+H⁺ = GSH-$NH_3$+H⁺ | $C_{10}H_{14}N_2O_6S+H^+$ | 291.0645 | 291.0652 | 0.67 |
| M-GSH+H⁺ = OPDA+H⁺ | $C_{18}H_{28}O_3+H^+$ | 293.2111 | 293.2116 | 0.48 |
| M-GSH-$H_2O$+H⁺ = OPDA-$H_2O$+H⁺ | $C_{18}H_{26}O_2+H^+$ | 275.2006 | 275.2011 | 0.54 |

**Figure S2:** Fragmentation pattern of GS-OPDA as determined by MSMS analysis. Peaks corresponding to each fragment in C) are assigned to by colour.

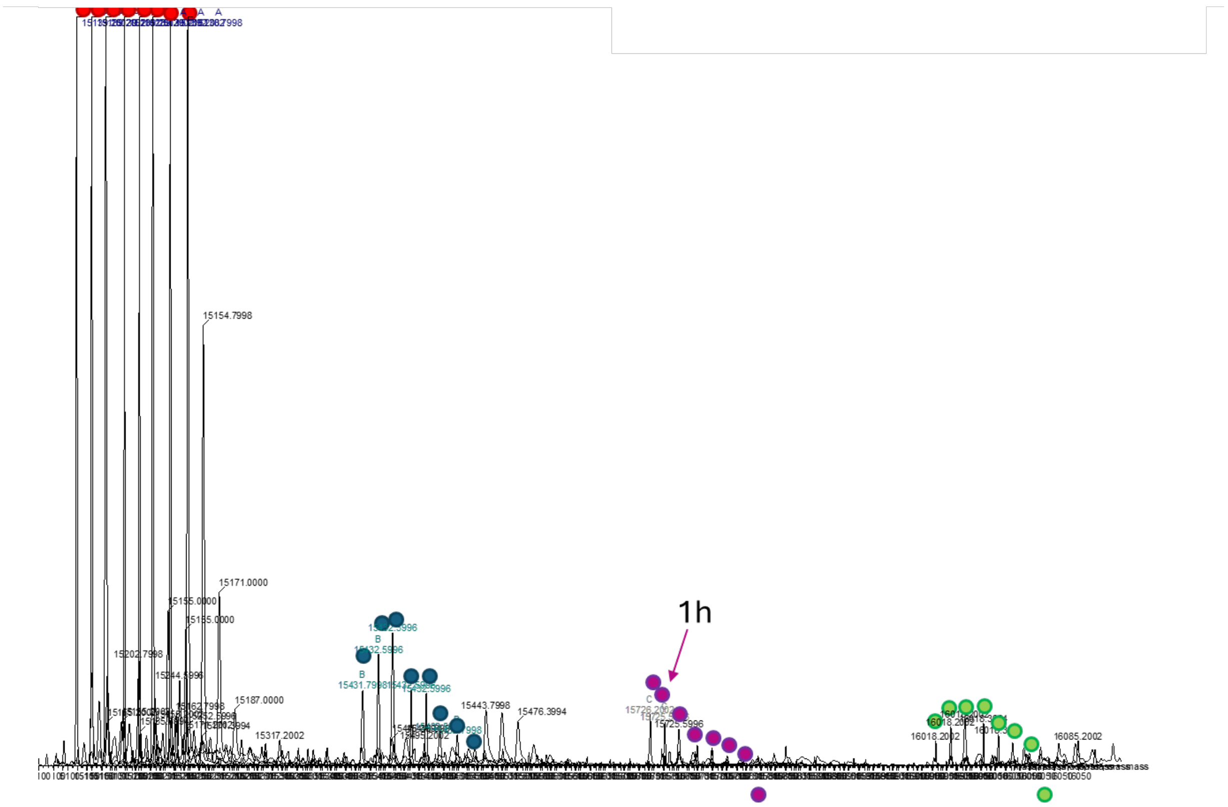


**Figure S3: MS analysis: GSH-triggered de-OPDAylation of TRX-h3.** Recombinant TRX was reduced, desalted, treated with OPDA for 16 h and desalted again. Samples were subjected to MS analysis at indicated time points.

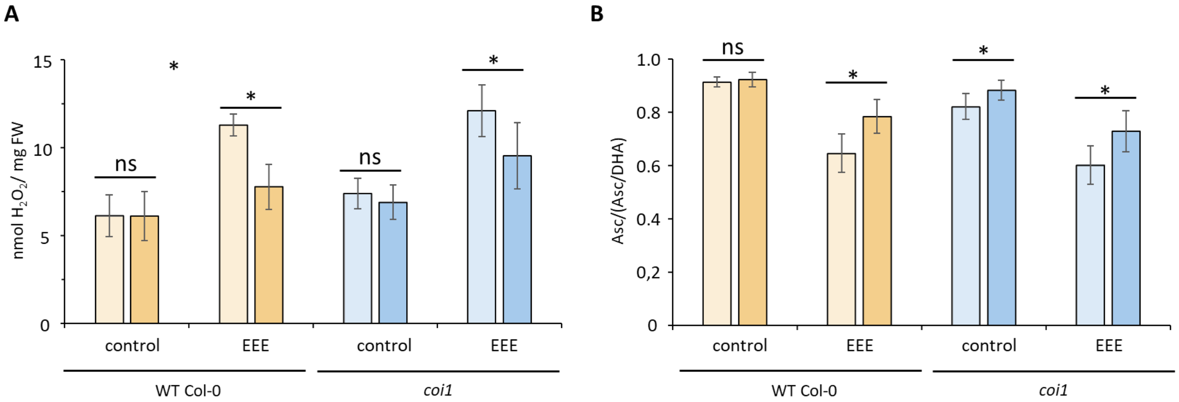


**Figure S4: Effect of OPDA on excess excitation energy (EEE)-induced $H_2O_2$ accumulation (A) and ratio of ascorbate (AsA) to total ascorbate pool (AsA + dehydroascorbate (DHA) (B).** Wild type plants were compared with *coi1*, defective in jasmonic acid signal perception. Half of the leaf discs were incubated in 25 µM OPDA solution while the other half were incubated in solution containing the same ethanol concentration for 16 h and subjected to EEE of 800 µmol photons·m$^{-2}$·s for 6 h. Light colors indicate control samples, dark colors indicate OPDA-treated samples. Asterisks denote significance of differences between groups that are treated and non-treated with OPDA ($p<0.05$; Student's t-test).

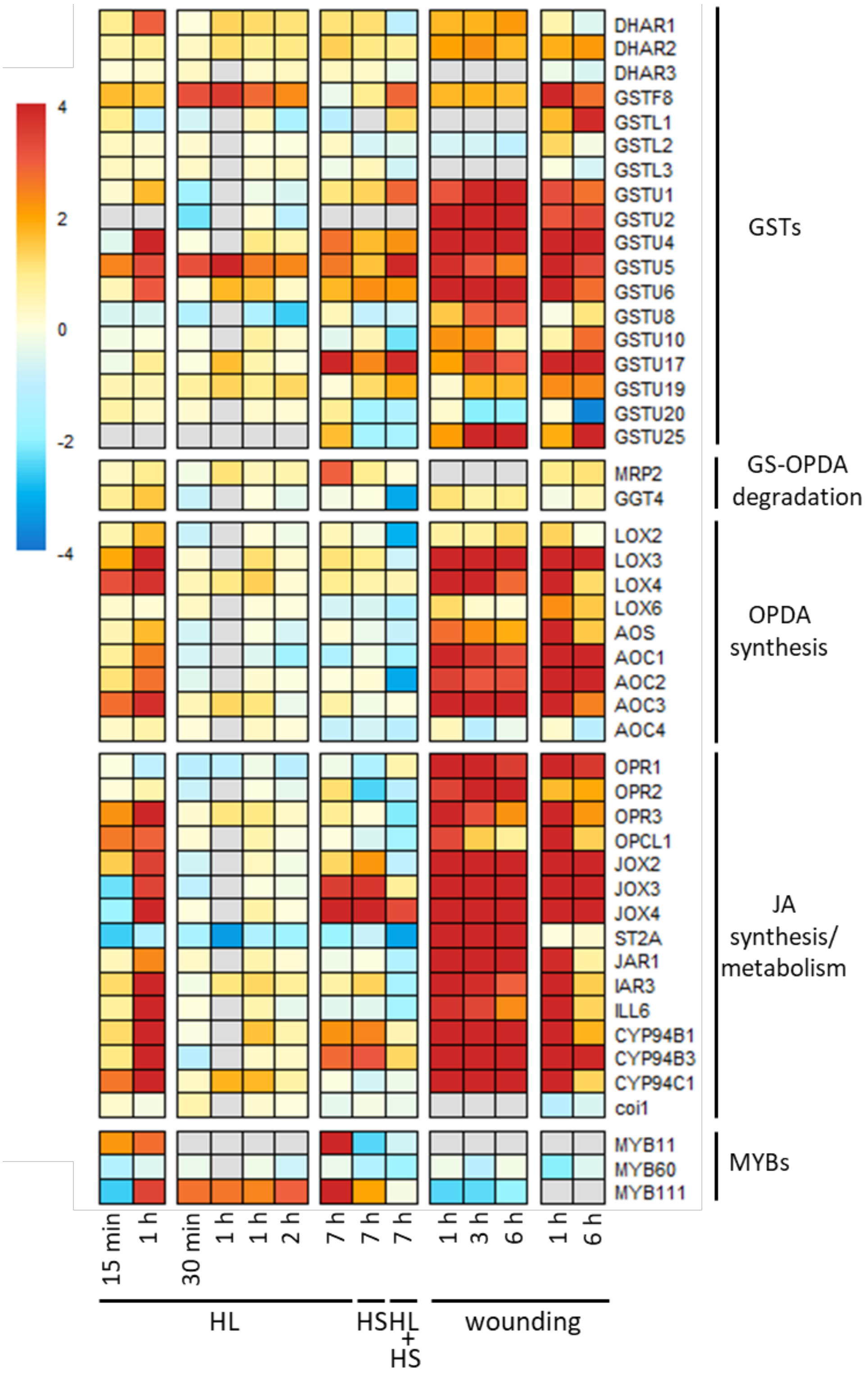


**Figure S3: Expression profiles of selected genes after stress treatment.** Transcriptome data was adapted by Kilic et al. (2025), Crisp et al. (2017), Balfagon et al., Xu et al. (2024) and Ikeuchi et al. (2017) (from left to right). HL=high light, HS=heat stress
